# Pervasive positive selection on X-linked ampliconic genes in primates

**DOI:** 10.64898/2026.03.09.710536

**Authors:** Emma Diepeveen, Meritxell Riera Bellés, Mikkel Heide Schierup

## Abstract

Mammalian sex chromosomes harbour ampliconic gene families, which are multi-copy genes with ≥97% sequence identity, predominantly expressed in testis tissue and essential for male fertility. Amplification of testis-specific genes is conserved across mammals, yet the specific gene families that expand show striking lineage-specific variation. Previous studies suggest a dynamic turnover with adaptive evolution for several of these families, but their analysis has been limited by the quality of reference genomes of repetitive regions. To characterize the molecular evolutionary processes of ampliconic gene families on both sex chromosomes, we analysed telomere-to-telomere genome assemblies from eight primate species spanning 25 million years of evolution. We identified 53 X-linked and 19 Y-linked ampliconic gene families with dynamic copy number variation. Gene conversion through palindromic pairing and tandem arrays maintained high sequence similarity despite accumulating mutations. X-linked families maintained conserved chromosomal positions despite copy number changes, while Y-linked families showed frequent positional turnover. Strikingly, multiple X-linked families (GAGE, SSX, CSAG, VCX) showed pervasive positive selection across the primate phylogeny and multiple (MAGEB, CT45, HSFX) showed lineage specific positive selection. Y-linked families predominantly evolve under purifying selection. These patterns could suggest that sperm competition, meiotic drive, or dosage-dependent selection drive the rapid, lineage-specific evolution of testis-expressed ampliconic genes in primates.

## Introduction

The mammalian X and Y chromosomes evolved from a pair of autosomes approximately 170 million years ago (Cortez et al. 2014). Following the emergence of the sex-determining locus on the proto-Y chromosome, a series of inversions suppressed XY recombination, across most of their length, excluding the pseudoautosomal regions (PARs). This resulted in great differentiation between the two chromosomes. The Y chromosome contains, besides the PAR and ancestral region (X-degenerate), long ampliconic regions with extensive intra-chromosomal homology. Ampliconic regions can be organized as tandem arrays as well as palindromes, long inverted repeats that undergo gene conversion, which counteracts the accumulation of deleterious mutations (Rozen et al. 2003; Skov et al. 2017; Hallast et al. 2023). Similarly, the X chromosome contains PAR(s), ancestral regions, and several palindromes (Miga et al. 2020; Makova et al. 2024). Within these regions, on both the X and the Y, lie ampliconic gene families with highly similar gene copies (≥97% sequence identity), and some families have expanded to high copy numbers. Most of these ampliconic genes are protein-coding and expressed exclusively in testis tissue, suggesting specialized functions in male gametogenesis (Mueller et al. 2013).

This pattern of testis-specific gene amplification is widespread across species with sex chromosomes, including humans, mice, and cattle (Soh et al. 2014; Hughes et al. 2020; Zhou et al. 2023). Studies in model organisms have revealed distinct evolutionary mechanisms underlying ampliconic expansion. In *Drosophila miranda*, which acquired neo-sex chromosomes through chromosomal fusion, new ampliconic genes arose through dramatic tandem amplification, with two apparent evolutionary drivers: testis-specific, dosage-sensitive genes amplified on the Y chromosome (potentially to increase male fertility), and meiosis-related genes co-amplified on both X and Y chromosomes (possibly reflecting genomic conflict over sex chromosome transmission) (Bachtrog et al. 2019). Similarly, the X- and Y-linked ampliconic genes *Slx* and *Sly* in mice exemplify co-amplification between sex chromosomes (Moretti et al. 2020; Arora and Dumont 2022; Kopania et al. 2022; Arlt et al. 2025). Functional studies demonstrate the importance of ampliconic genes for male fertility. Knockout of several ampliconic gene families results in infertility in mice, including RHOXF (Borgmann et al. 2016), SPACA5 (Park et al. 2026), TCP11 (Castaneda et al. 2020; TCP11 in mice is on chr7 not X/Y), and DMRTC1 (Kawamata and Nishimori 2006). In humans, deletions of X- and Y-linked ampliconic regions are associated with male infertility and spermatogenic failure, including SSX (C. Liu et al. 2023), H2BW (Bai et al. 2025), genes in AZFc regions such as DAZ, CDY and BPY (Xu and Pang 2022), and genes in the AFZa region, including RMBY and TSPY (Krausz and Casamonti 2017).

Despite these conserved patterns of testis-specific amplification and functional importance for male fertility across species, the specific gene families that undergo amplification show striking lineage-specific characteristics. Human and mouse X-ampliconic genes for example show little overlap; only 31% of human X-ampliconic genes have mouse orthologs, compared to 95% for single-copy genes. Yet ampliconic genes in both mice and humans show specialization in spermatogenesis (Mueller et al. 2013). This suggests rapid lineage-specific turnover of these gene families despite shared functions.

Most studies of ampliconic gene evolution have focused on the Y chromosome, due to their clinical role in male infertility. Recent studies of the Y chromosome have revealed substantial copy number variation within and between primate species, further supporting rapid evolutionary turnover in ampliconic gene families (Tomaszkiewicz et al. 2016; Skov et al. 2017; Ye et al. 2018; Lucotte et al. 2018; Vegesna et al. 2020). In contrast, X chromosome ampliconic genes have received comparatively less attention, with studies primarily examining copy number variation or positive selection in individual gene families. These studies provided important insights into rapid evolution: X-linked ampliconic families including GAGE (and its subfamilies PAGE and XAGE), CSAG, MAGE, and SSX showed signatures of positive selection in human-chimp comparisons (Stevenson et al. 2007; Y. Liu et al. 2008), and SPANX shows rapid evolution and amplification in great apes (Kouprina et al. 2004). However, the reference genome assemblies available at the time contained gaps and collapsed repetitive regions in ampliconic sequences, limiting the resolution of these regions and preventing comprehensive analysis of sequence evolution and copy number dynamics for both sex chromosomes across multiple primate lineages.

Recent telomere-to-telomere (T2T) assemblies of sex chromosomes in humans (Miga et al. 2020; Rhie et al. 2023), great apes (Makova et al. 2024), and crab-eating macaque (Zhang et al. 2025) have now resolved these previously intractable regions, enabling complete identification and annotation of ampliconic gene families across primates for the first time. While initial analyses have already described copy number variation for Y chromosome families (Makova et al. 2024), the opportunity now exists for comprehensive analysis of molecular evolution across multiple species for ampliconic genes on both sex chromosomes. Here we leverage these complete assemblies to examine the molecular evolution of X and Y ampliconic gene families across eight primate species, analysing copy number dynamics of ampliconic clusters, chromosomal gene movement, and the evidence of positive selection.

## Methods

### Detection of ampliconic gene families

To study ampliconic genes, we obtained telomere-to-telomere reference genomes from NCBI for six great ape species (human (*Homo sapiens),* bonobo (*Pan paniscus*), chimpanzee (*Pan troglodytes*), western lowland gorilla (*Gorilla gorilla*), Bornean orangutan (*Pongo pygmaeus*), and Sumatran orangutan (*Pongo abelii*)), one lesser ape (siamang gibbon (*Symphalangus syndactylus*)), and one monkey (crab-eating macaque (*Macaca fascicularis*)) (Miga et al. 2020; Rhie et al. 2023; Makova et al. 2024; Zhang et al. 2025). Multi-copy gene families were identified using the pipeline from Makova et al. 2024, based on the longest isoform protein sequences for all X and Y genes. Gene families were classified as multi-copy when ≥2 sequences showed >50% identity over >35% of their length with an e-value ≤0.001. To identify ampliconic gene families, we applied a ≥97% identity threshold (Supplementary Table 1). A gene family was considered ampliconic if at least one species contained ≥2 copies with ≥97% identity, based on a natural breakpoint in the pairwise sequence identity distribution for Y multi-copy genes (Makova et al. 2024). For each family, we compiled copy number counts, sequence class assignments across the chromosomes (as classified by Makova et al. 2024; Supplementary Table 2,3) for copy locations, and all pairwise identities of gene copies within each species.

To identify X-Y paralogous gene pairs (gametologs), we performed reciprocal blastp from BLAST+ v2.16.0+ searches between all X and Y chromosome protein sequences (Camacho et al. 2009). Paralogous pairs were retained if they showed ≥30% sequence identity over ≥50% of the alignment length with an e-value ≤0.001. Gene pairs were further classified based on their amplification status on each chromosome using the ampliconic classifications defined below to identify co-amplification of X and Y gene families.

### Clustering of ampliconic gene families

Nucleotide coding sequences were extracted for each gene in an ampliconic gene family. Isoforms were selected using the following priority hierarchy: RefSeq select tag if present; otherwise, no isoform tag; isoform X1; isoform 1 or a; or the lowest isoform number or alphabetically first isoform. This hierarchy was necessary because isoform naming was inconsistent within and across annotation files, and we wanted to use the first recorded isoform for each gene to remain consistent. It should be noted that transcript isoform diversity was not included in this study, even though this exists for at least some Y-linked ampliconic gene families (Tomaszkiewicz et al. 2023). We used blastn from BLAST+ v2.16.0+ to compare all sequences within an ampliconic family across species against a BLAST database built from the extracted sequences (Camacho et al. 2009). Genes were clustered into ampliconic clusters using transitive clustering (if genes A and B shared sequence identity and genes B and C shared sequence identity, genes A, B, and C formed a single cluster). Genes from different species belonged to the same ampliconic cluster if they showed ≥95% identity over 80% of their sequence length. For the macaque, we used a more lenient threshold of ≥90% identity over 80% of sequence length to account for its ∼5% divergence from the other species. Overall identity thresholds were lower than protein-based clustering because we used nucleotide sequences, and isoform length variation could affect clustering results. Individual gene copies in one species, forming its own cluster, were excluded from ampliconic cluster overview because they could not be compared.

### Identifying gene family movement across the chromosome

To assess gene family synteny across species, we first plotted all gene locations and orientations on the X and Y chromosomes for each species, coloured by sequence class (Makova et al. 2024). For this overview visualisation, we did not subset genes into ampliconic clusters to maintain clarity and reduce visual complexity. To visualise gene family movements across species, we created chromosome synteny plots using D3.js v6 (Bostock et al. 2011). Gene families were clustered based on genomic proximity using a 500 kb tolerance threshold, grouping nearby gene copies into single positional markers. For each gene family in each species, we calculated the average genomic position of clustered copies. When a gene family had multiple separated copy groups across a chromosome, each group was represented as a distinct cluster. Gene families were tracked across consecutive species in the phylogeny to visualise positional changes. For families with equal cluster numbers between adjacent species, we drew direct one-to-one connections. For families with unequal cluster numbers, we used nearest-neighbour matching for multi-cluster families (X chromosome: H2AB, MAGEB, PABPC, TMSB15B, and SPANX; Y chromosome: VCY, TSPY, RBMY, HSFY, DAZ, CDY, and BPY) to minimise visual complexity, connecting each cluster to its nearest genomic neighbour in the adjacent species. Other families with unequal cluster numbers were connected using all possible pairwise combinations to show potential splits or mergers. Centromeric regions were defined based on satellite annotations in the sequence classes and chromosomes were drawn to scale based on their actual lengths (Makova et al. 2024). Gene family labels were positioned on the left for families present in bonobo and on the right for families absent in bonobo, with automatic grouping of nearby labels to prevent overlap.

Clustering of genes near palindromes was assessed using permutation tests. For each species, we calculated the distance from each gene midpoint to the nearest palindrome boundary. To generate a null distribution, we randomly placed genes across the chromosome (uniform distribution from 0 to maximum chromosome coordinate) and calculated mean distances to palindromes (10,000 permutations). Statistical significance was assessed using permutation tests (p < 0.05). Clustering strength was quantified as the ratio of observed to expected mean distances.

### Species-specific positive selection tests

#### Pairwise positive selection tests

When at least one species had more than one gene in an ampliconic cluster, we performed codon-based alignment using MACSE v2.07 (-prog alignSequences); clusters containing only single genes per species were excluded from further analysis. Alignments were refined (-prog refineAlignment) and cleaned (-prog exportAlignment -codonForInternalStop NNN -codonForInternalFS charForRemainingFS ---) to enable downstream analyses (Ranwez et al. 2018). Pairwise synonymous (dS) and non-synonymous (dN) nucleotide substitution rates were calculated using the Modified Nei-Gojobori method with complete deletion of gaps in MEGA v11.0.13 (Kumar et al. 2012; Tamura et al. 2021). These values were averaged across all pairwise gene comparisons within each ampliconic cluster for every two-species comparison, yielding dN and dS estimates for all gene copies in each cluster across all species pairs. Some comparisons resulted in infinite *dN/dS* values when no synonymous substitutions were present, reflecting insufficient sequence divergence for reliable rate estimation. To test for significance, each ampliconic cluster alignment was bootstrapped 10,000 times by codon. For each bootstrap replicate, dN and dS were recalculated. We interpreted *dN/dS* as significantly greater than 1 (positive selection) if ≤0.05% of bootstrapped dN/dS values were below 1, and significantly less than 1 (purifying selection) if ≤0.05% of bootstrapped values were above 1.

#### Lineage-specific branch tests

Pairwise species comparisons of *dN/dS* values revealed that certain species showed positive selection more frequently than others across comparisons. Based on these patterns, we identified candidate species or clades that might be driving positive selection in each ampliconic cluster’s phylogeny. To test these hypotheses, we performed branch tests using PAML CODEML v.4.10.9 (Yang 2007), designating the candidate species or clades as foreground branches to test for lineage-specific positive selection. For all ampliconic gene clusters on the X and Y chromosomes, we used the multi-sequence codon alignments generated previously and randomly selected one gene per species from each alignment. This random selection was necessary because copy number variation between species within multi-sequence alignments could bias selection tests. Based on the selected genes, we pruned the unrooted species tree (((((bonobo, chimpanzee), human), gorilla), (S. orangutan, B. orangutan)), siamang, macaque)) to include only species present in each alignment, as some species lacked sufficiently similar copies to be included in certain ampliconic clusters. An unrooted tree was used because the branch models are time-reversible and do not assume a molecular clock, since all tests were done on clades that were not part of the root. All gaps were removed from alignments before running CODEML, and cleandata = 0 was used to preserve information in the data as recommended by Álvarez-Carretero et al. 2023. Alignments were converted from FASTA to PHYLIP format using FASTAtoPHYL.pl (Álvarez-Carretero et al. 2023). Using each alignment and pruned species tree, we ran PAML CODEML under the M0 (one-ratio) model and a branch model allowing ω to vary between foreground and background branches. We compared models using likelihood ratio tests (LRT), where the test statistic is defined as 2Δℓ = 2(ℓ1 − ℓ0), with ℓ0 and ℓ1 representing log-likelihoods under the null and alternative models, respectively. The LRT statistic follows a χ² distribution with degrees of freedom equal to the difference in free parameters between models. To assess robustness, we repeated this process six times for each ampliconic cluster with different randomly selected gene copies representing each species. This number was chosen to match the highest observed median cluster copy number (6.5 copies within one ampliconic cluster per species), ensuring that we could sample the majority of available gene copies and capture within-cluster sequence diversity.

### Positive site selection tests

To detect positive selection at codon sites beyond overall and species-specific ampliconic gene cluster estimates, we performed site tests using PAML CODEML v.4.10.9 (Yang 2007). Following the same approach as for branch tests, we randomly selected one gene per species from each multi-sequence codon alignment, cleaned the alignment of gaps, converted it to the phylip format and pruned the species tree accordingly, resulting in unrooted trees. Using each alignment and pruned species tree, we ran PAML CODEML under various site models that allow ω to vary across codons: the homogeneous M0 (one-ratio) model and the site-heterogeneous models M1a (nearly neutral), M2a (positive selection), M7 (beta), and M8 (beta & ω). We compared nested models (M0 vs M1a; M1a vs M2a; M7 vs M8) using likelihood ratio tests (LRT), as described above. This approach identified codons under positive selection. To assess robustness, we again repeated this process six times for each ampliconic cluster with different randomly selected gene copies representing each species. We excluded clusters with only 1-3 positively selected sites, as these likely represented false positives or alignment errors. We also excluded positively selected sites from clusters containing only two species, as the random sampling approach would base these results on only two sequences, limiting reliability. For clusters with positively selected sites, we mapped the sites to the corresponding positions in a human gene from that ampliconic cluster. We predicted protein domains using InterPro (Blum et al. 2025) and visualized the distribution of positively selected sites along the protein to assess for clustering patterns in specific protein domains.

### Concerted evolution of multicopy gene families

We performed four tests to assess the occurrence of gene conversion across copies within gene families. First, we calculated median GC3 content for coding sequences per gene family in each species and correlated these values with copy number. GC3 was calculated on the codon alignment made previously for *dN/dS* estimations. We expected gene families with higher copy numbers to show increased GC3 content due to GC-biased gene conversion because of increased opportunity for gene conversion (Galtier et al. 2001; Rousselle et al. 2019). The significance of each correlation slope was assessed by permuting the median GC3 versus copy number relationship across gene families. Second, we calculated pairwise sequence differences between all gene pairs within each ampliconic cluster per species. We used whole gene sequences (5’UTR, CDS exons, 3’ untranslated regions and introns) rather than coding regions alone to increase power, as gene conversion likely affects entire sequences. Whole gene sequences were retrieved per family per species and aligned using MAFFT v7.525 with 1000 iterations (Katoh and Standley 2013). After removing all gaps, we calculated pairwise differences and correlated them with palindrome status: both genes on opposite palindrome arms, both on the same arm, in different palindromes, one in a palindrome and one not, or both outside palindromes. Palindrome classifications were based on sequence classes from Makova et al. 2024. We expected genes on opposite palindrome arms to show greater similarity due to gene conversion. Third, we correlated pairwise differences within gene families with copy number, expecting larger families to show fewer pairwise differences due to increased opportunity for gene conversion. Fourth, we compared pairwise differences with both absolute genomic distance (base pairs calculated from midpoint to midpoint of the two genes). We expected gene conversion to decrease with distance, as it occurs more frequently in tandem repeat arrays, as demonstrated for TSPY and RBMY arrays on the Y chromosome (Hallast et al. 2023). The significance of these relationships was also assessed by permutation testing to account for non-independence of pairwise comparisons.

### Expression analysis during spermatogenesis

Single-nucleus and single-cell RNA-seq data analysed in this study were obtained from (Riera, M. 2025), including newly generated chimpanzee testicular snRNA-seq data from an individual housed at Copenhagen Zoo and previously published datasets for human, chimpanzee, bonobo, and rhesus macaque (Murat et al. 2023). Reads were aligned to the human reference genome (T2T-CHM13v2.0) using STAR/STARsolo, with UMI quantification allowing multi-mapping reads resolved by expectation–maximization (Nurk et al. 2022). Comparative analyses were restricted to 1:1 orthologous protein-coding genes and long non-coding RNAs identified by reciprocal best BLAST hits. Low-quality nuclei/cells, predicted doublets, ambient RNA contamination, and empty droplets were removed prior to downstream analysis. Data were normalized with Sctransform and integrated across species using scVI to generate a shared latent space (Lopez et al. 2018; Hafemeister and Satija 2019; Choudhary and Satija 2022). Cells were clustered with Leiden, and spermatogenic trajectories were inferred by diffusion pseudotime analysis, computed separately for diploid and haploid stages and rescaled for cross-species comparison. Cells were binned along X- and Y-bearing spermatid pseudotime trajectories, and log-normalized gene expression was modelled across pseudotime using a spline-based hierarchical Bayesian framework in PyMC (Abril-Pla et al. 2023). Expression was decomposed into shared (across-species) and species-specific components, and genes were classified as expressed or species-specific based on posterior credible intervals and log-fold change thresholds. All downstream analyses in the present study were performed using the processed, normalized, integrated, and pseudotime-resolved data as provided by the original publication.

## Results

### Characterization of the ampliconic gene family architecture across primates

Using telomere-to-telomere assemblies, we identified 53 ampliconic gene families on the X chromosome and 19 on the Y chromosome across all species examined (Figure 1a). All included primate species share the same evolutionary strata (Zhou et al. 2023), yet exhibited clade- and lineage-specific expansions in gene family copy number (Figure 1a,b). Of the X-linked families, 20 of 53 were ampliconic in all eight species (Figure 1a), while only 2 of 19 Y-linked families were ampliconic across species (Figure 1b). The proportion of clade-specific differences in status was higher on the Y chromosome (13) than on the X (7) (p=2.48×10⁻⁵, binomial test), reflecting the broad conservation of X-linked gene families in a single-copy, multi-copy or ampliconic state. Of the Y-linked families, five were present in more than half of the species analysed (TSPY, CDY, DAZ, HSFY, RBMY). We additionally detected species-specific expansions: 6 X-linked and 6 Y-linked families were ampliconic in only a single species. Some of these families were entirely restricted to that one species: on the X chromosome, collagen alpha-1(IV) chain-like was found only in human; on the Y chromosome, KRT18Y was specific to bonobo, MTRNR2-like 17 to chimpanzee, centriole and centriolar satellite protein OFD1-like to gorilla, and TAF11L2 to siamang (Figure 1a,b).

**Fig 1.**
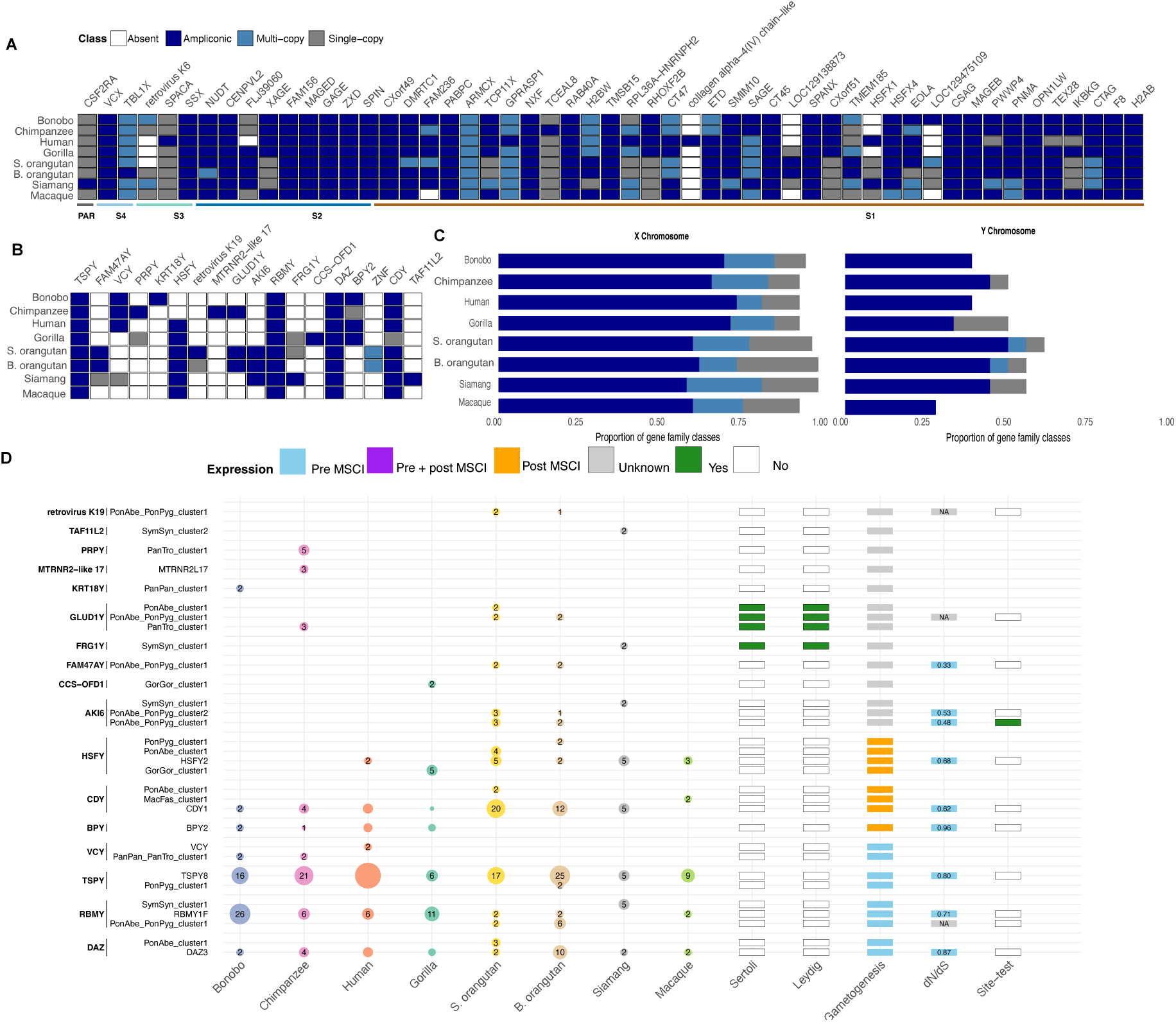
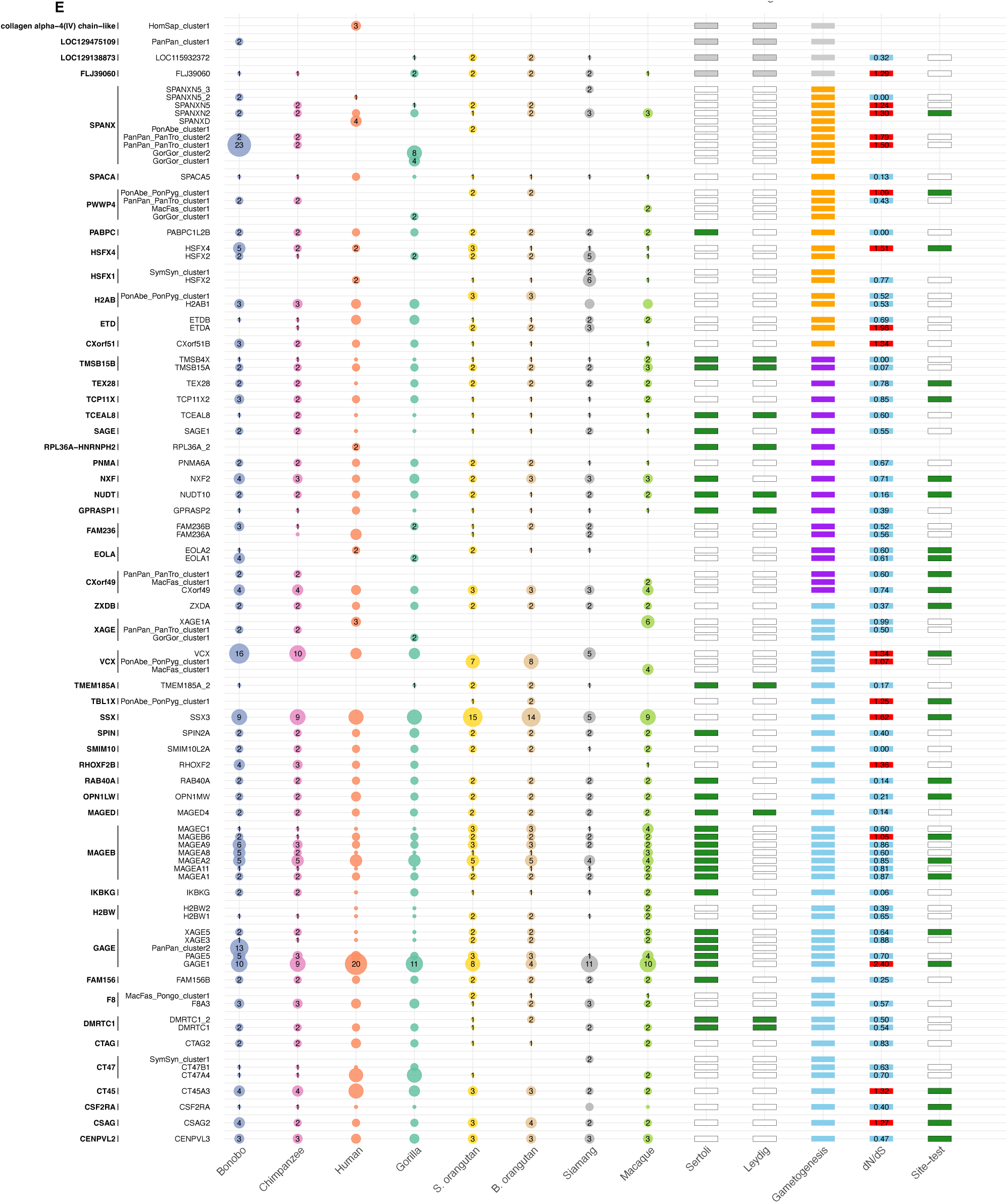
Evolution of ampliconic gene families on the X and Y chromosomes. **a, b.** Gene family comparison across primate species (**a**: X chromosome; **b**: Y chromosome). Gene families vary in copy number across species, being ampliconic (≥97% identity) in some species and multi-copy (50-97% identity) or single-copy in others. Families are ordered by human chromosomal coordinates and grouped by evolutionary strata. **c.** Proportion of gene families in each copy number category across species (left: X; right: Y). **d, e.** Ampliconic cluster structure and evolution (**d**: Y; **e**: X). Multi-copy gene families are divided into ampliconic clusters: gene copies sharing ≥97% identity with each other but <97% identity with other family members form separate clusters. Species-specific cluster names (e.g., PonAbe_PonPyg_cluster1) indicate clusters restricted to those species (Species abbreviations: PanPan (bonobo), PanTro (chimpanzee), HomSap (human), GorGor (gorilla), PonAbe (Sumatran orangutan), PonPyg (Bornean orangutan), SymSyn (siamang), MacFas (macaque)). Circle size and numbers indicate copy number per cluster; absence indicates no sufficiently similar copies. Expression annotations: green blocks indicate expression in testis somatic cells (Sertoli/Leydig); spermatogenesis timing shows expression relative to MSCI (pre-MSCI, post-MSCI, both, or unknown). Selection metrics: *dN/dS* values from PAML phylogenies (blue <1, indicating purifying selection; red ≥1 indicates possible positive selection); green blocks in the site-test column indicate positively selected sites detected in an ampliconic cluster.

Many ampliconic gene families belong to broader multi-copy gene families, defined as having copies with 50-97% sequence identity (compared to ≥97% for ampliconic genes). Within these multi-copy families, the most similar copies (≥97% identity) form discrete ampliconic clusters. For example, the MAGE family is a large multi-copy family containing multiple distinct ampliconic clusters (Figure 1c). This clustering pattern is consistent with the idea that some multi-copy families represent diverged remnants of ancestral ampliconic expansions. In some cases, different species lineages have independent ampliconic clusters within the same multi-copy family. For instance, two orangutan species share a VCX ampliconic cluster (PonAbe-PonPyg-Cluster1; Figure 1e) distinct from the VCX cluster found in other primates, indicating that orangutan VCX copies have diverged (below 97% identity) from other primate VCX copies to form a separate ampliconic cluster, while still belonging to the broader VCX multi-copy family.

Gene families with higher total copy numbers showed greater copy number variation across species on both the X and Y chromosomes (Supplementary Figure 1), as previously observed within human and great apes (Lucotte et al. 2018; Ye et al. 2018; Vegesna et al. 2020). This pattern is consistent with the increased probability of copy-number change at high copy number (Ghenu et al. 2016). Notably, cancer-testis families (13 on the X and 2 on the Y) were amongst the most variable families in copy number (Supplementary Figure 1). The X-linked cancer-testis families have been previously described as fast evolving (Stevenson et al. 2007). Y chromosome families showed particularly high copy numbers and wider variation than most X chromosome families: TSPY reached 44 copies in human, CDY 20 copies in B. orangutan and RBMY 26 copies in bonobo, an order or magnitude higher than most X chromosome families.

Some gene families showed consistent amplification across all species, while others displayed lineage-specific expansion patterns. Of the larger gene families, GAGE and MAGEA2 were consistently amplified across species, though bonobo possessed an additional GAGE ampliconic cluster, absent in all other species. SSX showed consistent amplification except for additional expansion in orangutans. In contrast, several gene families show pronounced amplification in only one or a few species: SPANX, GAGE, VCX, and RMBY exhibited extreme amplification in bonobo, while CT45 and CT47 show higher copy number in gorilla and human than in other species. TSPY displayed the widest amplitude of variation, ranging from pronounced amplification in humans, intermediate levels in bonobo, chimpanzee and orangutan and substantially lower copy numbers in gorilla siamang and macaque (Figure 1c,d)

Across primates, we identified ampliconic gametolog pairs using reciprocal BLAST searches (≥30% sequence identity) and filtering for ampliconic status on both chromosomes. There was no enrichment in ampliconic gametologs compared to non-ampliconic gametologs (Supplementary Table 4). This revealed two commonly co-amplified gene families across species: HSFX/Y and VCX/Y, although both HSFY and VCX were independently lost in some species. Several additional paralogous gene pairs showed amplification on one of the sex chromosomes: TSPY/X (Y-ampliconic) and RBMY/X (Y-ampliconic, X-multicopy), TBL1X/Y (X-ampliconic), TMSB/TMSB4Y (X-ampliconic). CSF2RA, which resides in the pseudoautosomal region, was present on both chromosomes but expanded only in siamang on the X.

#### Expression timing of ampliconic genes

Most ampliconic gene families are expressed in the testis (Supplementary Table 5, Figure 1d, e). Expression pattern during spermatogenesis may reveal functional constraints and selective pressures: genes expressed before meiotic sex chromosome inactivation (MSCI) must maintain function during early germ cell development, while genes expressed after MSCI escape silencing and may be subject to different selective forces, including potential meiotic drive. Expression in somatic testis cells (Sertoli and Leydig cells) versus germ cells may indicate distinct functional roles. Using single-cell RNA-seq data from human, chimpanzee, bonobo and (rhesus) macaque (Murat et al. 2023; see methods), we quantified expression across cell types and gametogenic stages. For germ cells, we categorized expression timing relative to meiotic sex chromosome inactivation (MSCI): pre-MSCI expression (undifferentiated and different spermatogonia, and leptotene through pachytene spermatocytes) or post-MSCI expression (round and elongated spermatids).

Some X-chromosome gene families show expression during both pre-and post-MSCI phases (Figure 1e), whereas Y-chromosome families show expression exclusively in either pre-MSCI or post-MSCI phases (Figure 1d). Families with post-MSCI expression show minimal expression in testis somatic cells. Of the pre-MSCI expressed genes, for both the X- and Y linked, expression is highest in differentiated spermatogonia and leptotene spermatocytes, after which they are silenced. This same trend is seen for genes expressed pre- and post-MSCI, where most have an expression peak in differentiated spermatogonia and leptotene spermatocytes and are reactivated in round spermatids after meiotic silencing. Expression data are unavailable for a subset of families (10 for the Y-linked and 3 for the X linked), because these families are not present in humans.

Paralogous gene pairs show divergent expression patterns. HSFY was expressed only post-MSCI and primarily in human and macaque but not in bonobo or chimpanzee (in which we also did not detect any gene copies), while its X-linked paralog HSFX was expressed post-MSCI across all species. VCY shows expression in human both pre- and post-MSCI, while VCX was expressed throughout pre- and post-MSCI phases in all species and to a higher level than VCY. This lower expression of VCY than VCX had been described in humans before (Vegesna et al. 2019). RBMY (Y-ampliconic) and RBMX (X-multicopy) were both expressed pre-MSCI across species, with RBMY showing particularly high expression in bonobo and human, possibly corresponding to extreme amplification in bonobo (Supplementary Figure 2).

### Synteny of ampliconic gene families

We next examined the genomic distribution of gene families by grouping genes into proximity blocks, as new copies typically arise through tandem duplication or within nearby palindromes (Supplementary Figures 5, 6). When tracing the synteny of these blocks across species, we found that X-linked gene families have conserved chromosomal positions (Figure 2). Only four X chromosome families showed distal gain or loss of gene copies while retaining their primary proximity blocks: TBL1X, SPANX, TCEAL8 and RPL36A-HNRNPH2 (Supplementary Figure 3). SPANX was a notable exception, having lost/not gained an entire block in Macaque. Many gene families had at least some copies within palindromic regions (36/53 X-linked families). The distribution of X-linked gene families closely traced locations of these palindromes, with genes significantly closer to palindromes than expected by chance in all seven species with palindrome annotations (permutation test, all p < 0.001; Supplementary Figure 4). The degree of clustering (observed/expected distance) was similar across species (clustering ratio = 0.36 ± 0.06, range: 0.30-0.47), with genes positioned approximately 2-3x closer to palindromes than expected under random distribution.

**Fig 2.**
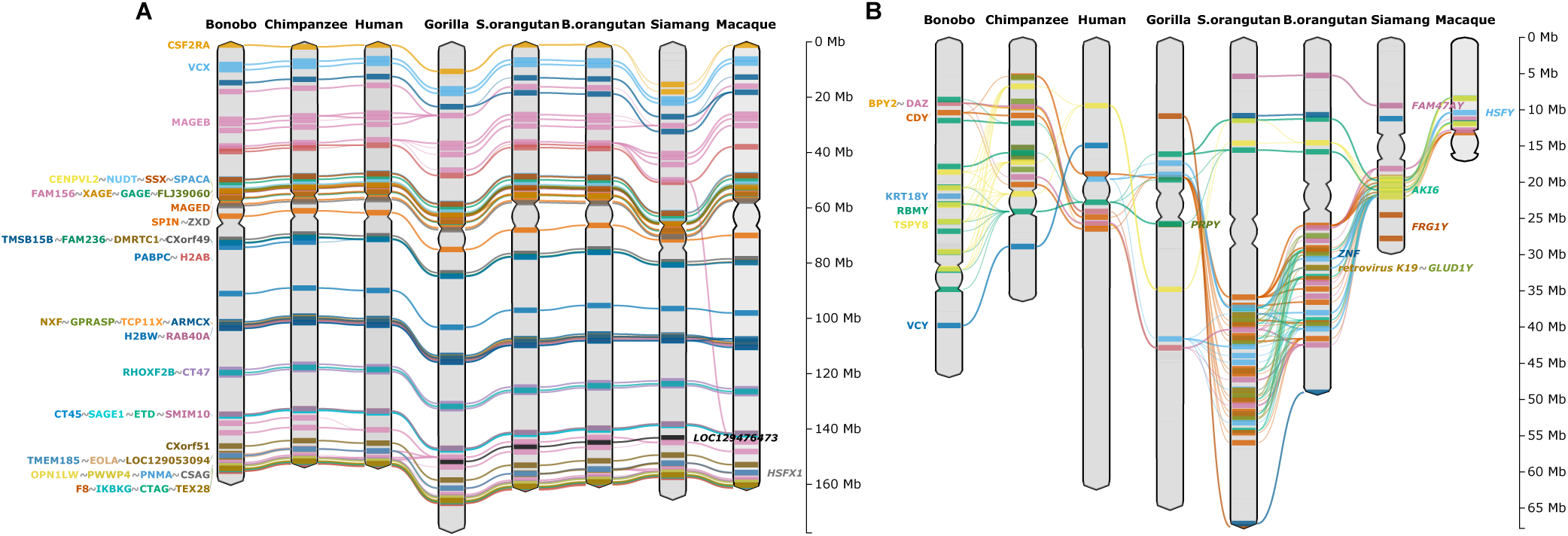
Gene family synteny across primate species. **A**: X chromosome; **B**: Y chromosome. Gene families on the X chromosome maintain conserved positions across species, while Y chromosome gene families show greater positional variation. Each gene family is represented by its average chromosomal position; families with multiple separated groups of gene copies are shown as multiple blocks. Gene family labels present in the bonobo are positioned on the left and on the right for families absent in the bonobo, with automatic grouping of nearby labels to prevent overlap.

By contrast, Y-linked gene families displayed marked positional turnover, including recurrent gain and loss events across lineages (Figures 1a; Figure 2; Supplementary Figure 6). This increased variability is consistent with the overall sequence class composition of the Y chromosome, which shows much greater variation across species compared to the X chromosome (Makova et al. 2024; Supplementary Figure 5, 6). Similar to the patterns observed on the X chromosome, many Y-linked gene families were associated with palindromic regions (10/17 gene families). Y-linked gene copies also showed significant clustering near palindromes in most species (6/7 species, p < 0.05, not for the S. orangutan, Supplementary Figure 4), though with greater variability across the phylogeny (clustering ratio = 0.46 ± 0.23, range: 0.21-0.89) compared to the X chromosome. Some great apes showed the strongest clustering (gorilla, human, orangutan: ratios 0.21-0.34, approximately 3-5x closer than expected), while the S. orangutan showed less concentration of gene families around palindromes (ratio = 0.89). Although this is likely because palindromes are spread across the whole chromosome in general in the S. orangutan (Supplementary Figure 6).

#### Gene conversion homogenises ampliconic genes within a family

We subsequently tested whether gene conversion homogenises sequences among ampliconic gene copies. The ampliconic families occur in two main structural contexts: within palindromes or sequentially in tandem repeats. We hypothesized that gene conversion homogenisation would be strongest for copies located on opposite arms of palindromes, where arm-to-arm gene conversion has been extensively characterized. The inverted-repeat structure of the palindromes facilitates pairing, which in turn promotes gene conversion between copies, providing a mechanism for purging deleterious mutations and maintaining sequence integrity in ampliconic genes (Figure 3a; Rozen et al. 2003; Skov et al. 2017; Hallast et al. 2023). Both the X and Y chromosomes harbour multiple palindromes and as described above, many gene families occur within or near these palindromes (Supplementary Figures 4, 5, 6).

**Fig 3.**
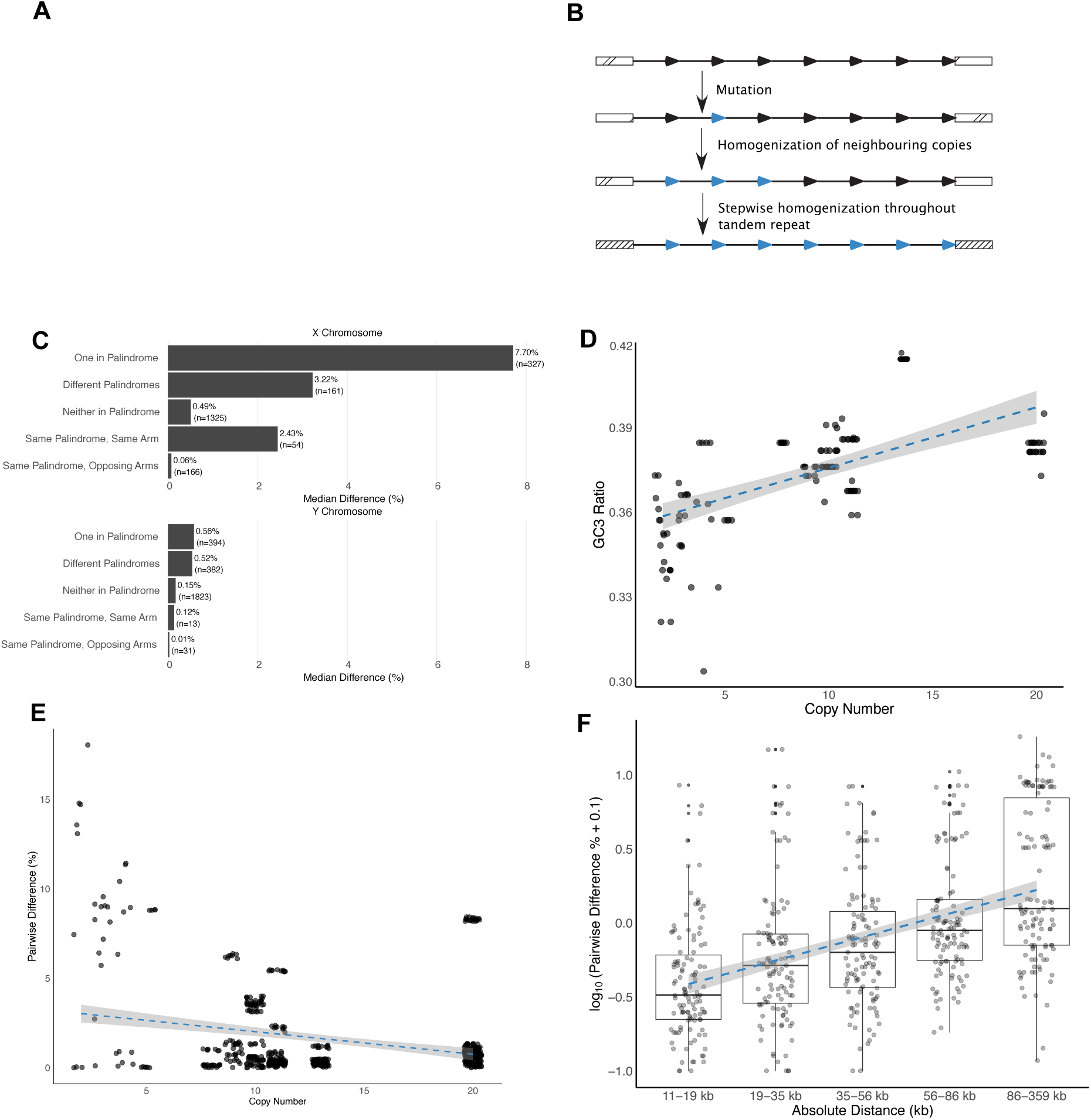
Gene conversion in ampliconic gene families. **a.** Schematic of palindrome structure promoting arm-to-arm pairing. **b.** Schematic of gene conversion homogenisation in tandem arrays**. c**. Pairwise sequence differences between gene copies grouped by palindrome classification (top: X chromosome, 2.033 comparisons in 49 families; bottom: Y chromosome, 2.643 comparisons in 17 families, across 7 species). Genes on opposite palindrome arms show the highest sequence similarity. Y chromosome genes show greater overall similarity than X chromosome genes regardless of palindrome status. **d-f**. Gene conversion signals in the X-linked GAGE gene family. **d.** GC3 content increases with copy number (135 genes across 8 species). **e.** Pairwise sequence differences decrease with copy number (575 pairwise comparisons across 8 species). **f.** Physically closer gene pairs show greater sequence similarity. Pairwise differences can appear slightly negative due to log(x + 0.1) transformation used for visualisation.

We compared full gene sequences (including non-coding regions) for all pairwise combinations of gene copies within each ampliconic cluster per species. We then classified each gene pair based on whether genes co-occurred in palindromes and, if so, whether they were on the same or opposite palindrome arms. As expected, genes occurring on opposing arms of a palindrome show the highest sequence similarity (0.06% median difference on the X chromosome (CI: 0.027-0.107%) and 0.01% on the Y chromosome (CI:0-0.059%) (Figure 3c). Overall sequence differences were much lower on the Y chromosome than on the X chromosome (with the highest median of 0.56% difference on the Y chromosome versus 7.70% as the highest on the X chromosome). Interestingly, gene copies that both did not occur in palindromes also showed low pairwise differences (0.15% on the Y (CI:0.15-0.16%) and 0.49% on the X (CI: 0.43-0.61%)). These copies were in tandem repeats, suggesting that gene conversion also operates within tandem arrays to some extent.

We tested for the possibility of homogenization across gene copies within tandem repeats, using the GAGE family, which has high copy numbers and occurs primarily in tandem repeats rather than palindromes (Supplementary Figure 5; Supplementary Table 2,6). Three independent lines of evidence supported gene conversion in tandem arrays. First, median GC3 content of coding regions increased with copy number (slope = 0.003, p = 0.0048), consistent with GC-biased gene conversion (gBGC) (Rousselle et al. 2019; Figure 3d). This pattern is expected because more copies provide a greater opportunity for gene conversion. Second, pairwise differences across whole gene sequences decreased with copy number (slope = −0.688, p < 0.001), indicating more homogenization with higher copy number (Figure 3e). Third, gene copies in closer physical proximity showed higher sequence similarity than distant copies (slope = 2.48×10⁻⁵, p < 0.001) (Figure 3f), consistent with proximity-dependent gene conversion across tandem repeats. This homogenization pattern resembles that previously reported for TSPY protein-coding copies on the Y chromosome, where recombination between palindrome arms and/or direct repeats maintains high sequence similarity (Bonito et al. 2021; Makova et al. 2024). This suggests that, in addition to gene conversion occurring on the Y, it also occurs on the X chromosome, keeping ampliconic copies similar.

### Pervasive positive selection in Gene families on multiple levels

#### Positive selection at the species level

To test for positive selection, we used PAML’s maximum likelihood framework for estimating *dN/dS* rates (ω) across primate evolutionary history by integrating over all gene sequences in the context of a phylogeny (Yang 2007). Across species, we detected no signs of positive selection across the entire phylogeny for any of the Y chromosome families, consistent with previous findings (Makova et al. 2024; Figure 1d). However, pairwise *dN/dS* estimates revealed lineage-specific selection patterns. DAZ showed elevated *dN/dS* in comparisons between human and both bonobo and chimpanzee, and branch tests showed significantly higher ω in the hominin clade (ω=∼2-3 versus ∼0.7), suggesting accelerated evolution on these lineages (Figure 4b). RBMY showed elevated *dN/dS* (∼4) in comparisons between the chimpanzee and bonobo. All other Y chromosome families showed patterns consistent with purifying selection or neutral evolution (Supplementary Table 7).

**Fig 4.**
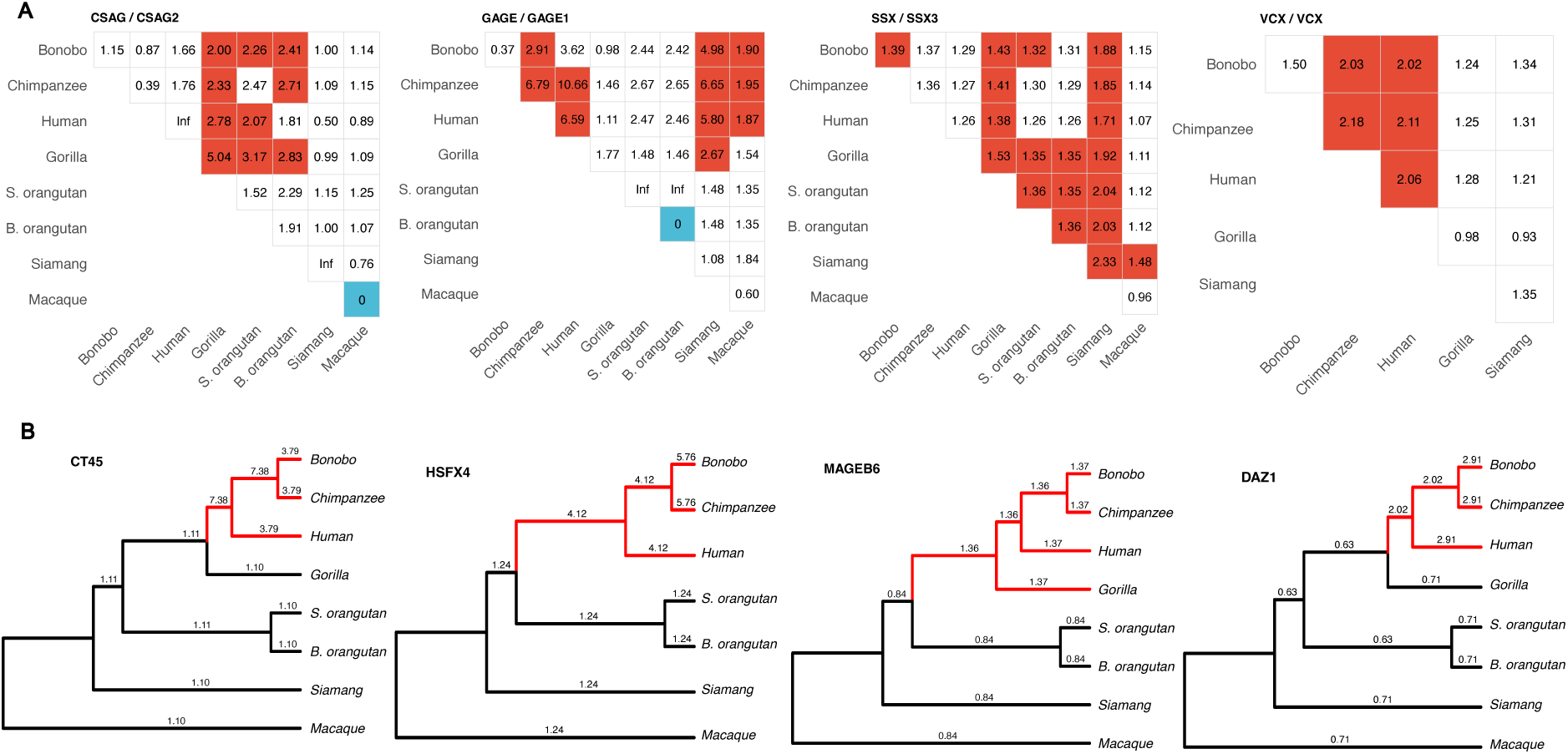
Species-specific positive selection in gene families. **a.** Pairwise *dN/dS* estimates across gene copies in ampliconic clusters. SSX, GAGE, CSAG, and VCX show elevated evolutionary rates across multiple lineages, with no single species or clade driving the signal, suggesting pervasive positive selection across the phylogeny. Red colouring indicates significant *dN/dS* > 1; Blue colouring indicates significant *dN/dS <1,* both after bootstrapping. **b**. PAML branch test results. Omega (ω) values are shown for each branch on the gene tree, with foreground branches in red. Likelihood ratio tests (LRT) compared null and branch models. PAML detected significant positive selection in the hominini clade (chimpanzee, bonobo, and human) for X-linked ampliconic clusters CT45 and HSFX4 and the Y-linked DAZ cluster. The ampliconic cluster MAGEB6 shows positive selection in the African great ape clade (chimpanzee, bonobo, human, and gorilla).

In contrast to the Y chromosome, several X-linked gene families showed evidence of positive selection across the species tree (Supplementary Table 8). To assess whether these ω values were elevated compared to other X chromosome genes, we compared them to 50 randomly selected X-linked genes, which had a median ω of 0.16 (IQR: 0.08-0.31). The gene families under positive selection were exclusively expressed in testis, either before or after meiotic sex chromosome inactivation (MSCI). Post-MSCI genes showed a higher proportion with evidence of positive selection (8/16, 50%) compared to pre-MSCI genes (9/36, 25%), though this difference was not statistically significant (p=0.15, proportional binomial test) (Figure 1e). This pattern is consistent with findings that expression evolution is fastest for post-meiotic cell types (Murat et al. 2023).

To identify which lineages or clades drove positive selection patterns, we estimated pairwise *dN/dS* values across species. Focusing on the largest ampliconic cluster within a gene family, five gene families (CSAG, GAGE, SSX, VCX, MAGE) show higher *dN/dS* values across nearly all pairwise comparisons (Figure 4a), suggesting pervasive rather than lineage-specific positive selection. We estimated higher *dN/dS* values for VCX genes in hominins, CSAG in gorilla and orangutans, and FLJ390960 (MAGED4) primarily in orangutan lineages. However, these patterns were not supported by branch tests (Supplementary table 9), likely due to short sequence lengths and uniformly elevated *dN/dS* values across the phylogeny, both of which reduce the power of branch-specific tests (Duchemin et al. 2023). Signals of positive selection on these gene families have been previously reported, though not across this broad of species sampling (Stevenson et al. 2007; Y. Liu et al. 2008). Within the MAGE family, both ampliconic and single-copy clusters showed evidence of positive selection, particularly in (African) great apes (Figure 4b; Supplementary Table 8). Several families also showed elevated *dN/dS* when comparing ampliconic paralogs within a species, particularly the rapidly evolving families CSAG, GAGE, SSX and VCX (Figure 4a; Supplementary Table 8).

In addition to these gene families showing pervasive selection, we also detected evidence of positive selection in the hominini clade for CT45 and HSFX3/4, with ω estimates 3- to 4-fold higher along these branches, respectively (CT45: 3.79-7.38 in hominini versus 1.10; HSFX3/4 4.12-5.67 in hominini versus 1.24) (Figure 4b). The function of CT45 is unknown but interestingly, humans have twice as many copies as the chimpanzee and bonobo of CT45 (Figure 1e). HSFX is predicted to function as a DNA transcription regulator of stress response genes (Hästbacka et al. 2025). The expression of HSFX3/4 is slightly higher in the hominini, than in macaque (Supplementary Figure 2). MAGEB6 exhibited positive selection in the African great ape clade, with ω estimates approximately 0.5 higher than all other branches (Figure 4b).

#### Positive selection at the site level

We next used PAML to search for evidence of positive selection at a subset of amino acids within the gene families (site-specific tests) (Yang 2007). Interestingly, several gene families showed positively selected sites despite exhibiting overall neutral or purifying selection across the species tree (Supplementary note 1; Supplementary table 10; Supplementary Figure 7). These families may harbour positively selected sites amid broader neutral or purifying selection. There was no positive selection for sites found in families on the Y chromosome (Figure 1d; Supplementary Table 11). For two rapidly evolving gene families on the X (GAGE and CT45A) (Supplementary table 10), we detected 1-3 sporadic positively selected sites, and these sites did not cluster in specific protein domains or regions. In other rapidly evolving families (SSX, MAGEB6, VCX), positively selected sites were generally dispersed across protein sequences. VCX showed a compact cluster of positively selected sites located outside its known VCX/Y domain (Figure 5a). MAGEB6 displayed dispersed positive selected sites throughout the protein, with an apparent hotspot in the first helix of its MAGE domain (Figure 5c). The SSX cluster showed a scattering of positively selected sites with a notable exception: the SSXRD domain contained no positively selected sites, while the KRAB domain accumulated eight positive sites (Figure 5b). These rapidly evolving gene families and many other ampliconic gene families encode primarily intrinsically disordered proteins (Figure 5; Supplementary figure 7), lacking stable 3D structures. Disordered regions are essential for cellular processes including transcriptional control, cell signalling, and subcellular organization (Holehouse and Kragelund 2024). Intrinsically disordered regions are frequent targets of positive selection in humans and across the mammalian phylogeny, thought to be hotspots for genetic innovation (Afanasyeva et al. 2018).

**Figure 5.**
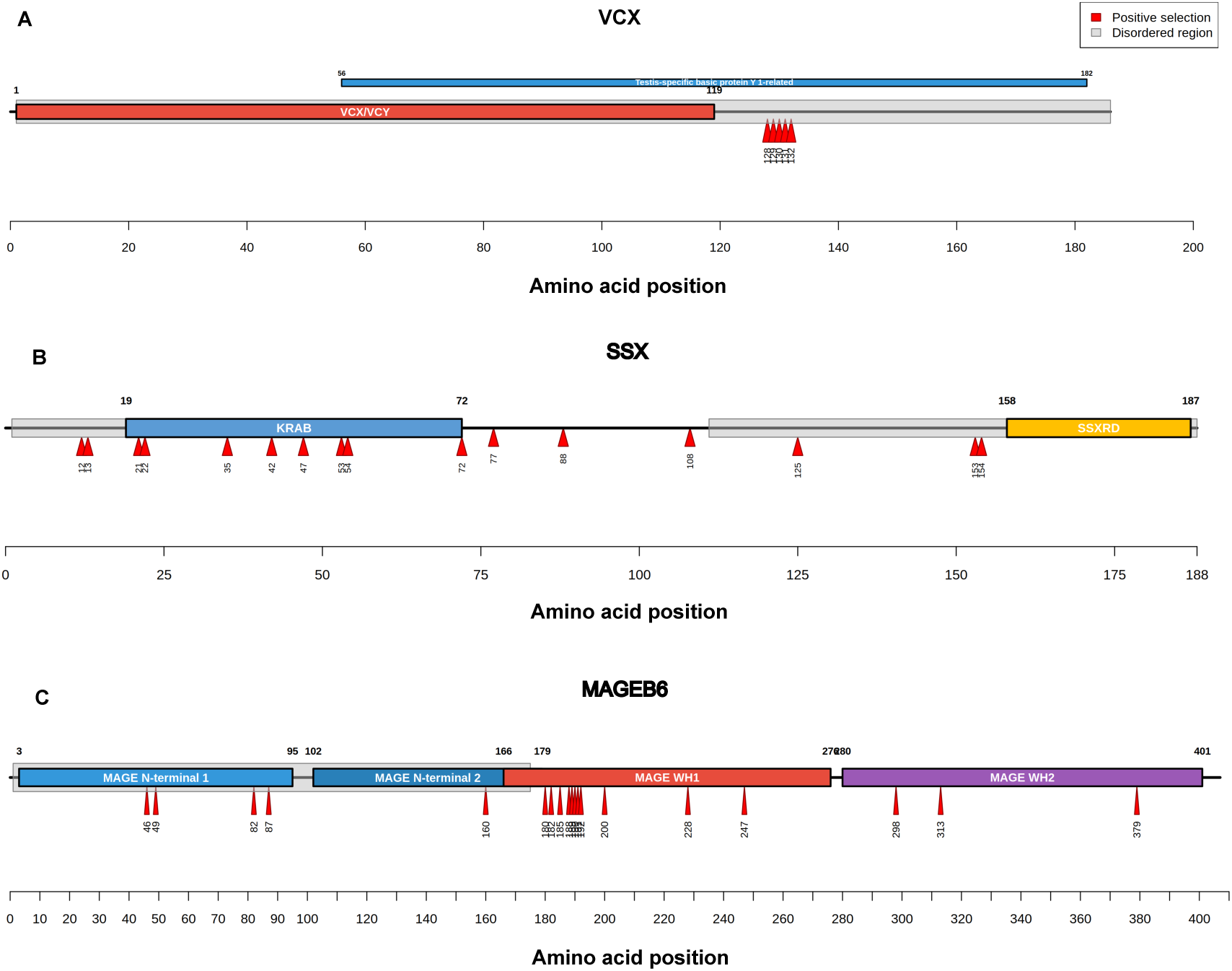
Distribution of positively selected sites in rapidly evolving gene families. Protein domains are based on human amino acid sequences as predicted by InterPro (Blum et al., 2025). Positively selected sites were identified by PAML site tests. **a.** VCX shows a compact cluster of positively selected sites in the intrinsically disordered region outside the known VCX/Y domain. **b.** SSX shows positively selected sites dispersed across the protein sequence, with absence in the SSXRD domain and concentration in the KRAB domain. **c.** MAGEB6 shows positively selected sites dispersed throughout the protein, with a hotspot in the first helix of the MAGE domain.

## Discussion

Using complete telomere-to-telomere sex chromosome assemblies across eight primate species, we found pervasive positive selection on X-linked ampliconic genes, with several families showing elevated *dN/dS* ratios across the primate phylogeny. In striking contrast, Y-linked families predominantly evolved under purifying selection. Gene conversion operates through palindromic structures and tandem arrays to homogenize sequences within families, maintaining high sequence similarity despite accumulating mutations. Copy number variation (CNV) on X- and Y-linked families is extensive, with some families showing extreme lineage-specific amplifications, yet X-linked families maintain conserved chromosomal positions across species despite these dramatic copy number changes, while Y-linked families show frequent positional turnover. The connection between gametogenesis-specific expression, positive selection and extreme CNV of ampliconic families suggest strong selective pressures are acting on these genes. Several non-mutually exclusive mechanisms may explain the rapid evolution.

First, sperm competition intensity varies with mating systems and could influence ampliconic gene evolution since their expression is primarily in the testis. Species with multi-male mating systems (polyandry and polygynandry) experience higher sperm competition, evidenced by increased relative testis size and increased meiotic index (Skakkebæk et al. 2022). The elevated evolutionary rates or increased copy number in some ampliconic families could reflect adaptation to species-specific sperm competition regimes. Consistent with this hypothesis, bonobo (the species experiencing the highest sperm competition among those examined) shows extreme copy number expansion in several families including GAGE, SPANX, VCX, and RBMY. The functional significance of such copy number variation is supported by findings in humans that RBMY copy number correlates with sperm count and motility (Yan et al. 2017), suggesting direct phenotypic consequences of ampliconic gene dosage and therefore possibly selection in bonobo. However, the relationship between copy number and sperm competition is not universal, nor does high copy number appear universally advantageous. Some families show extreme amplification in species with lower sperm competition: TSPY reached 44 copies in human (low sperm competition species) and CDY 20 copies in B. orangutan (intermediate sperm competition species). In humans, selection acts against nonreference copy numbers for many ampliconic families, suggesting possible optimal copy number ranges (Lucotte et al. 2023). TSPY is a notable exception as it shows high copy numbers across all species, with most copies occurring outside palindromes, contrasting with the typical palindrome- associated amplification pattern. This together suggests that additional selective forces beyond sperm competition are at play.

Second, meiotic drive may contribute to ampliconic gene evolution, where co-amplified X and Y genes act as drivers and suppressors in genomic conflict over sex chromosome transmission, which could contribute to ampliconic gene evolution. Previous studies in mouse and cattle have shown co-amplification of paralogous gene families on the X and Y chromosomes (Soh et al. 2014; Hughes et al. 2020). In mice, knockout of either Slx (X-linked) or Sly (Y-linked) in mice causes transmission distortion and non-Mendelian sex chromosome segregation (Moretti et al. 2020; Arora and Dumont 2022; Kopania et al. 2022; Arlt et al. 2025). This co amplification reflects an evolutionary arms race, where amplification of a driver selects for either suppression or counter-drive. We confirmed co-amplification in two paralogous pairs (VCX/Y and HSFX/Y), consistent with a similar type of potential meiotic drive. If meiotic drive operates in primates, we would expect stronger effects in species with lower sperm competition, where relaxed selection on sperm quality permits transmission bias. Expression patterns might support this prediction: VCY is expressed in human and HSFY is expressed in human and macaque (low/intermediate sperm competition) but both not in bonobo and chimpanzee (high sperm competition). However, their X-linked counterparts VCX and HSFX are expressed across all species. The expression timing of the gene pairs differs: HSFX/Y is expressed post-MSCI in round and elongated spermatids while VCX/Y is expressed pre-MSCI in leptotene spermatocytes. Notably, loss of HSFY is associated with male infertility in humans (Shinka et al. 2004), suggesting functional importance despite species-specific expression patterns.

Meiotic drive need not involve paralogous pairs directly. In *Drosophila*, selfish distorters exploit a meiotic checkpoint gene, to cause segregation distortion by eliminating competing sperm (Ridges et al. 2026). Similar mechanisms could operate in primates, with different gene families competing through amplification, though such effects are difficult to detect.

A third mechanism for possible amplification is dosage compensation, which has been documented in mice and *Drosophila* (Lucchesi and Kuroda 2015) and may operate in primate ampliconic genes. Dosage compensation could provide an alternative explanation for the VCX/Y and HSFX/Y patterns we observed. Rather than representing meiotic drive through genomic conflict, continued expression of VCX and HSFX across all species could compensate for loss of VCY and HSFY expression (Vegesna et al. 2019). This interpretation is consistent with our finding that VCX and HSFX maintain expression across all species despite variation in their Y-counterpart expression. Beyond compensation, another possible mechanism is dosage regulation, where ampliconic genes show lower expression than their non-ampliconic homologs. In great apes, some Y ampliconic gene families maintain stable expression levels despite substantial copy number variation (Vegesna et al. 2020). In humans specifically, most Y ampliconic gene families show expression levels lower to their non-Y homologs, indicating this dosage regulation (Vegesna et al. 2019). This down-regulated expression could reflect sub-functionalization (e.g., testis-specific expression that benefits germline development) or relaxed selection. The latter is supported by the finding that Y ampliconic genes show weaker purifying selection (higher *dN/dS*) than non-ampliconic Y genes (Betrán et al. 2012). This relaxed constraint, combined with gene conversion, could facilitate rapid functional differentiation (Vegesna et al. 2019; Makova et al. 2024).

These mechanisms possibly driving ampliconic gene evolution, sperm competition, meiotic drive and dosage compensation, are not mutually exclusive and likely interact. Distinguishing among these alternatives will require additional data and approaches. Our analysis used expression data from four species; broader sampling across all eight species and incorporation of intraspecific variation will be crucial for understanding these dynamics. Intraspecific CNV is substantial for at least several ampliconic families. In humans, the X-linked families GAGE, CT45, and SPANX show extensive copy number variation, as do the Y-linked families TSPY, DAZ, BPY, CDY and HSFY, often associated with palindrome duplications (Lucotte et al. 2018). Families with extensive copy number variation across humans, have been shown to have significantly higher expression in pachytene spermatocytes during meiosis (Lucotte et al. 2018). Using finer-resolution cell-stage data, we found that families with high copy numbers and described interspecific copy number variation (GAGE, SSX, VCX, MAGE, CT45, CSAG) are predominantly expressed in differentiated spermatogonia and leptotene spermatocytes (pre-MSCI). These are also families that show positive selection across primates. Overall, it seems like there is less high CNV in post-MSCI expressed families, but they do show signals of selection. Integrating expression patterns, copy number variation, and positive selection across both inter- and intraspecific scales will ultimately reveal which evolutionary mechanisms drive the strange dynamics of ampliconic gene families.

## Code Availability

Code used in the manuscript is available at GitHub link: https://github.com/emmadiepeveen/Ampliconic-Evolution-Primates

