## Supplementary Figures for "Pervasive positive selection on X-linked ampliconic genes in primates"

### Supplementary Materials

#### Supplementary Tables

*See separate Additional File 2.xlsx file*

**Supplementary Table 1**: X- and Y ampliconic gene family pairwise identities per species
**Supplementary Table 2**: X-linked ampliconic gene family information
**Supplementary Table 3:** Y-linked ampliconic gene family information
**Supplementary Table 4:** Enrichment analysis of gametologs among X ampliconic genes
**Supplementary Table 5:** Gene functions and literature summary of XY ampliconic families
**Supplementary Table 6**: Slope estimates for gene conversion tests
**Supplementary Table 7**: Pairwise *dN/dS* estimates per ampliconic gene cluster on the Y
**Supplementary Table 8**: Pairwise *dN/dS* estimates per ampliconic gene cluster on the X
**Supplementary Table 9**: PAML branch test results for ampliconic gene clusters
**Supplementary Table 10:** PAML site test results per X-linked ampliconic gene cluster
**Supplementary Table 11:** PAML site test results per Y-linked ampliconic gene cluster

#### Supplementary Figures

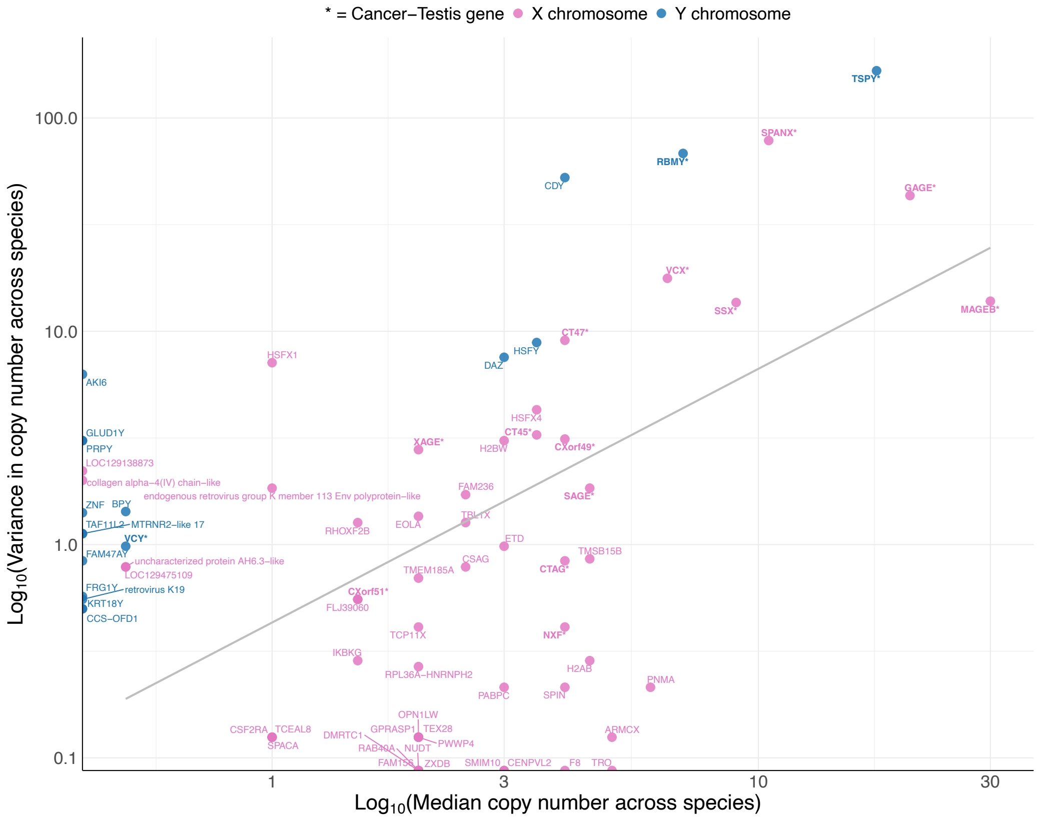


**Supplementary Fig 1. Relationship between gene family size and copy number variability across species.** Median and variance of copy number were calculated for each gene family across all eight species. The grey line shows the line of best fit. Cancer-testis genes are indicated in bold with asterisks (*). Pink labels indicate X chromosome families; blue labels indicate Y chromosome families.

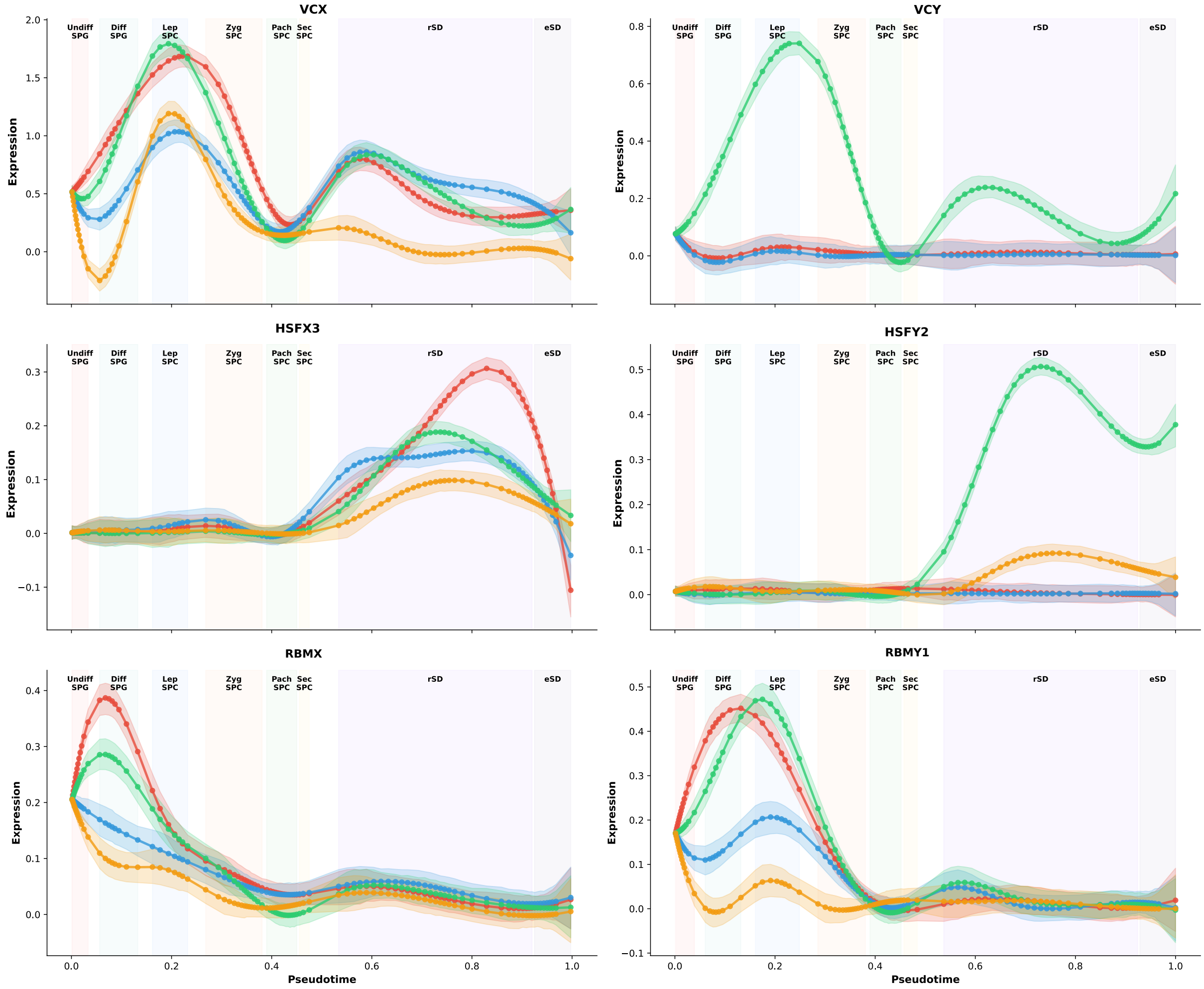


**Supplementary Fig 2. Expression of paralogous multicopy genes across pseudotime of spermatogenesis.** Trajectories show log-normalized gene expression levels across pseudotime-ordered spermatogenic cell stages for X- and Y-linked paralogous gene pairs. Undiff SPG = Undifferentiated spermatogonia; Diff SPG = Differentiated spermatogonia; Lep SPC = Leptotene spermatocytes; Zyg SPC = Zygotene spermatocytes; Pach SPC = Pachytene spermatocytes; Sec SPC = Secondary spermatocytes; rSD = Round spermatids; eSD = Elongated spermatids. Red = bonobo, blue = chimpanzee, green = human, orange = macaque.

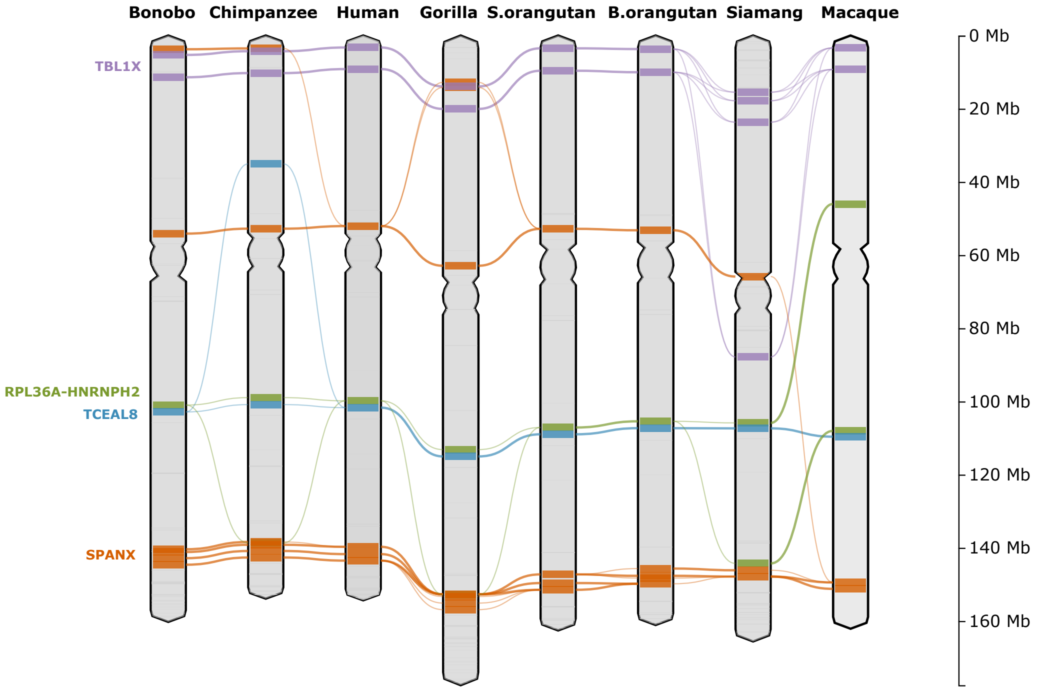


**Supplementary Fig 3. Positional variation of four X chromosome gene families across species.** Each gene family is represented by its average chromosomal position; families with multiple separated copy groups are shown as multiple blocks. Movement of these X-linked families are characterized by appearance or loss of copies at distant chromosomal locations rather than complete translocation of the entire gene family.

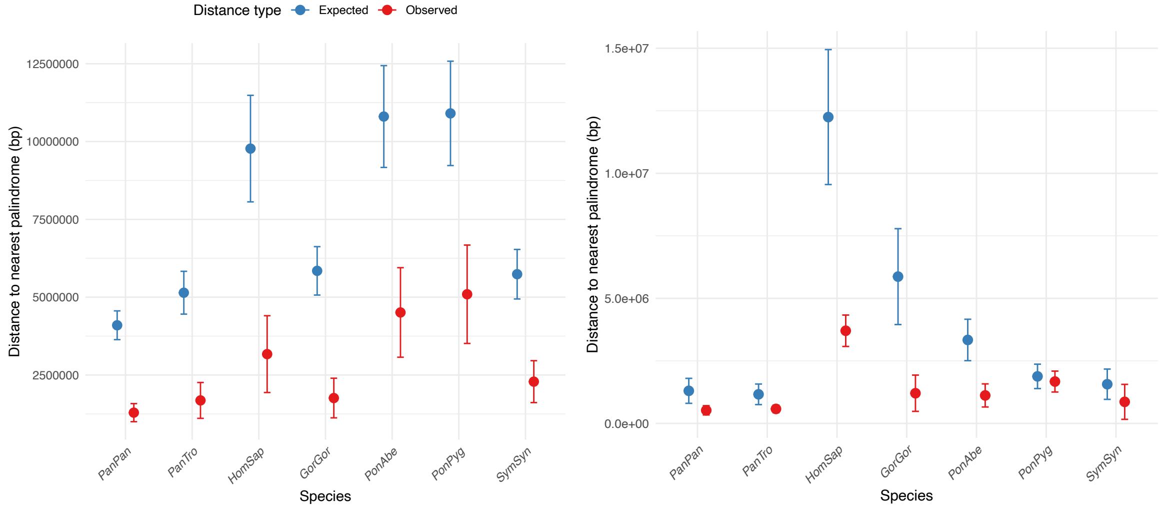


**Supplementary Fig 4. Observed versus expected distance to palindromes on the X (left) and Y (right) chromosomes**. Points show mean observed (red) and expected (blue) distances to the nearest palindrome, with 95% confidence intervals. Expected distances were derived from 10,000 permutations of random gene placement across the chromosome. Ampliconic gene families concentrate significantly closer to palindromes than expected by chance in most species. Species abbreviations: PanPan (bonobo), PanTro (chimpanzee), HomSap (human), GorGor (gorilla), PonAbe (Sumatran orangutan), PonPyg (Bornean orangutan), SymSyn (siamang). Palindrome annotations from Makova et al. 2024; not available for macaque.

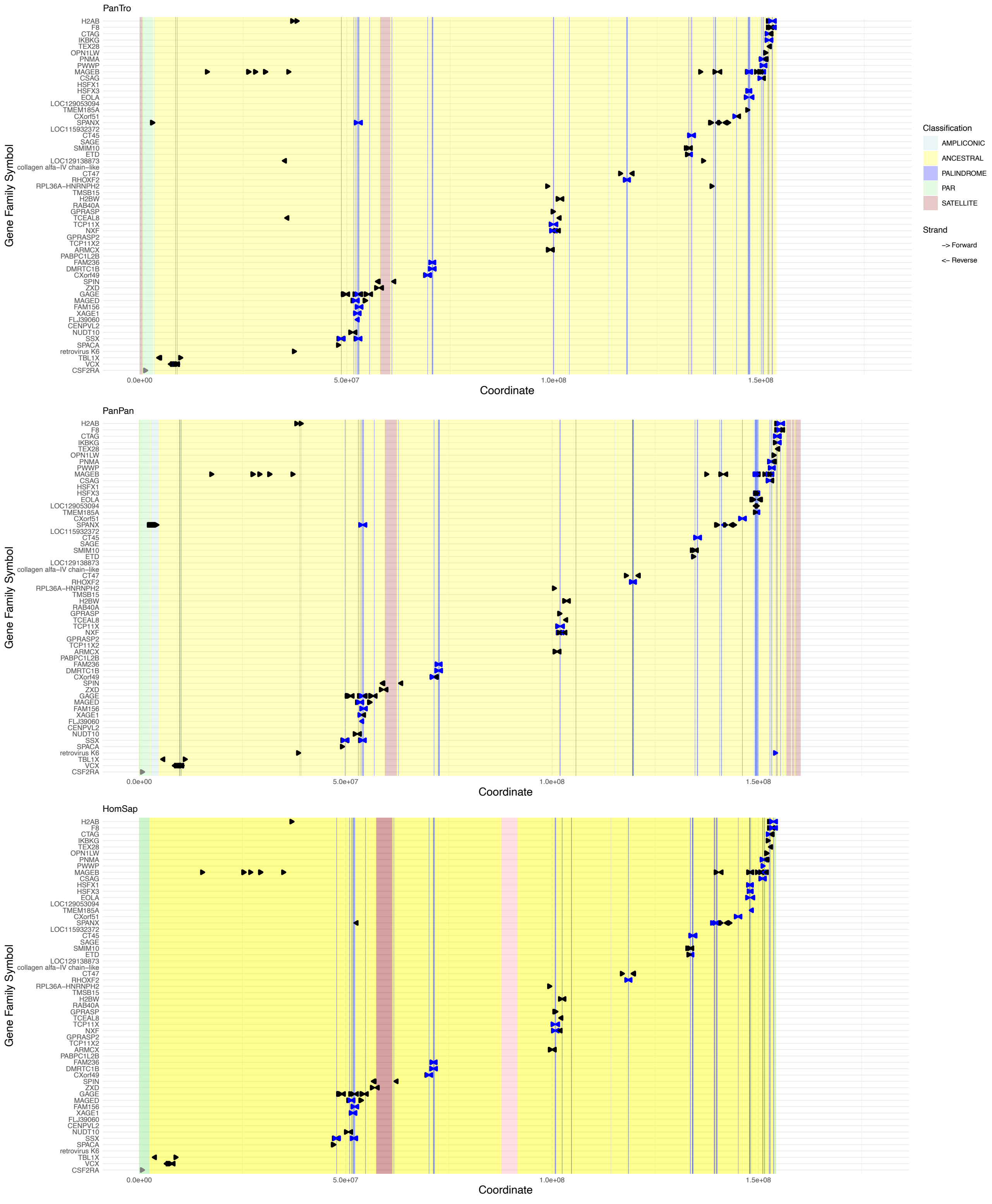
PanTro
Gene Family Symbol
H2AB
F8
CTAG
IKBKG
TEX28
OPN1LW
PNMA
PWWP
MAGEB
CSAG
HSFX1
HSFX3
EOLA
LOC129053094
TMEM185A
CXorf51
SPANX
LOC115932372
CT45
SAGE
SMIM10
ETD
LOC129138873
collagen alfa−IV chain−like
CT47
RHOXF2
RPL36A−HNRNPH2
TMSB15
H2BW
RAB40A
GPRASP
TCEAL8
TCP11X
NXF
GPRASP2
TCP11X2
ARMCX
PABPC1L2B
FAM236
DMRTC1B
CXorf49
SPIN
ZXD
GAGE
MAGED
FAM156
XAGE1
FLJ39060
CENPVL2
NUDT10
SSX
SPACA
retrovirus K6
TBL1X
VCX
CSF2RA
0.0e+00
5.0e+07
1.0e+08
1.5e+08
Coordinate
PanPan
Gene Family Symbol
H2AB
F8
CTAG
IKBKG
TEX28
OPN1LW
PNMA
PWWP
MAGEB
CSAG
HSFX1
HSFX3
EOLA
LOC129053094
TMEM185A
CXorf51
SPANX
LOC115932372
CT45
SAGE
SMIM10
ETD
LOC129138873
collagen alfa−IV chain−like
CT47
RHOXF2
RPL36A−HNRNPH2
TMSB15
H2BW
RAB40A
GPRASP
TCEAL8
TCP11X
NXF
GPRASP2
TCP11X2
ARMCX
PABPC1L2B
FAM236
DMRTC1B
CXorf49
SPIN
ZXD
GAGE
MAGED
FAM156
XAGE1
FLJ39060
CENPVL2
NUDT10
SSX
SPACA
retrovirus K6
TBL1X
VCX
CSF2RA
0.0e+00
5.0e+07
1.0e+08
1.5e+08
Coordinate
HomSap
Gene Family Symbol
H2AB
F8
CTAG
IKBKG
TEX28
OPN1LW
PNMA
PWWP
MAGEB
CSAG
HSFX1
HSFX3
EOLA
LOC129053094
TMEM185A
CXorf51
SPANX
LOC115932372
CT45
SAGE
SMIM10
ETD
LOC129138873
collagen alfa−IV chain−like
CT47
RHOXF2
RPL36A−HNRNPH2
TMSB15
H2BW
RAB40A
GPRASP
TCEAL8
TCP11X
NXF
GPRASP2
TCP11X2
ARMCX
PABPC1L2B
FAM236
DMRTC1B
CXorf49
SPIN
ZXD
GAGE
MAGED
FAM156
XAGE1
FLJ39060
CENPVL2
NUDT10
SSX
SPACA
retrovirus K6
TBL1X
VCX
CSF2RA
0.0e+00
5.0e+07
1.0e+08
1.5e+08
Coordinate
Classification
AMPLICONIC
ANCESTRAL
PALINDROME
PAR
SATELLITE
Strand
−> Forward
<− Reverse

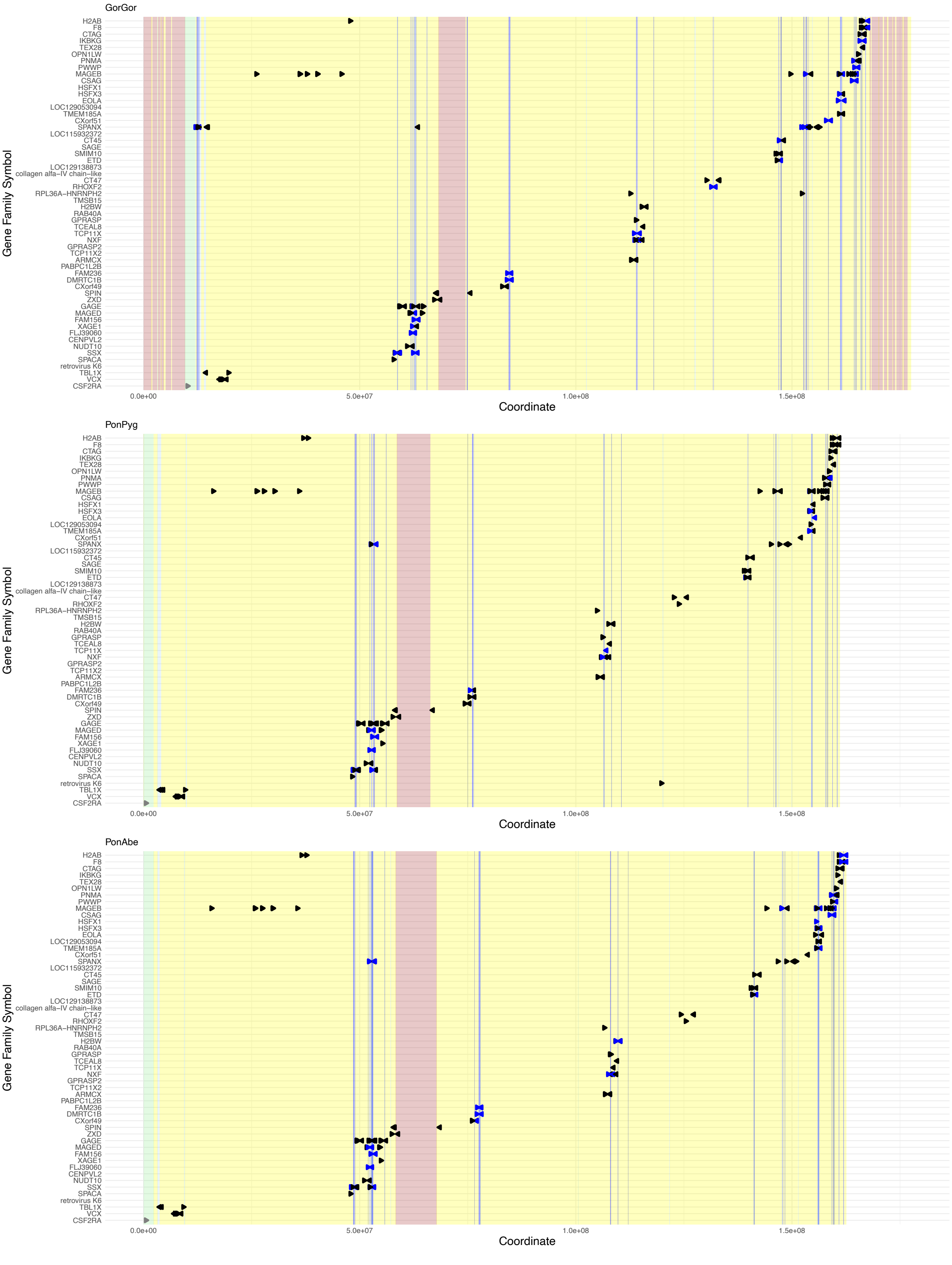

GorGor
Gene Family Symbol
H2AB
F8
CTAG
IKBKG
TEX28
OPN1LW
PNMA
PWWP
MAGEB
CSAG
HSFX1
HSFX3
EOLA
LOC129053094
TMEM185A
CXorf51
SPANX
LOC115932372
CT45
SAGE
SMIM10
ETD
LOC129138873
collagen alfa−IV chain−like
CT47
RHOXF2
RPL36A−HNRNPH2
TMSB15
H2BW
RAB40A
GPRASP
TCEAL8
TCP11X
NXF
GPRASP2
TCP11X2
ARMCX
PABPC1L2B
FAM236
DMRTC1B
CXorf49
SPIN
ZXD
GAGE
MAGED
FAM156
XAGE1
FLJ39060
CENPVL2
NUDT10
SSX
SPACA
retrovirus K6
TBL1X
VCX
CSF2RA
0.0e+00
5.0e+07
1.0e+08
1.5e+08
Coordinate
PonPyg
Gene Family Symbol
0.0e+00
5.0e+07
1.0e+08
1.5e+08
Coordinate
PonAbe
Gene Family Symbol
0.0e+00
5.0e+07
1.0e+08
1.5e+08
Coordinate

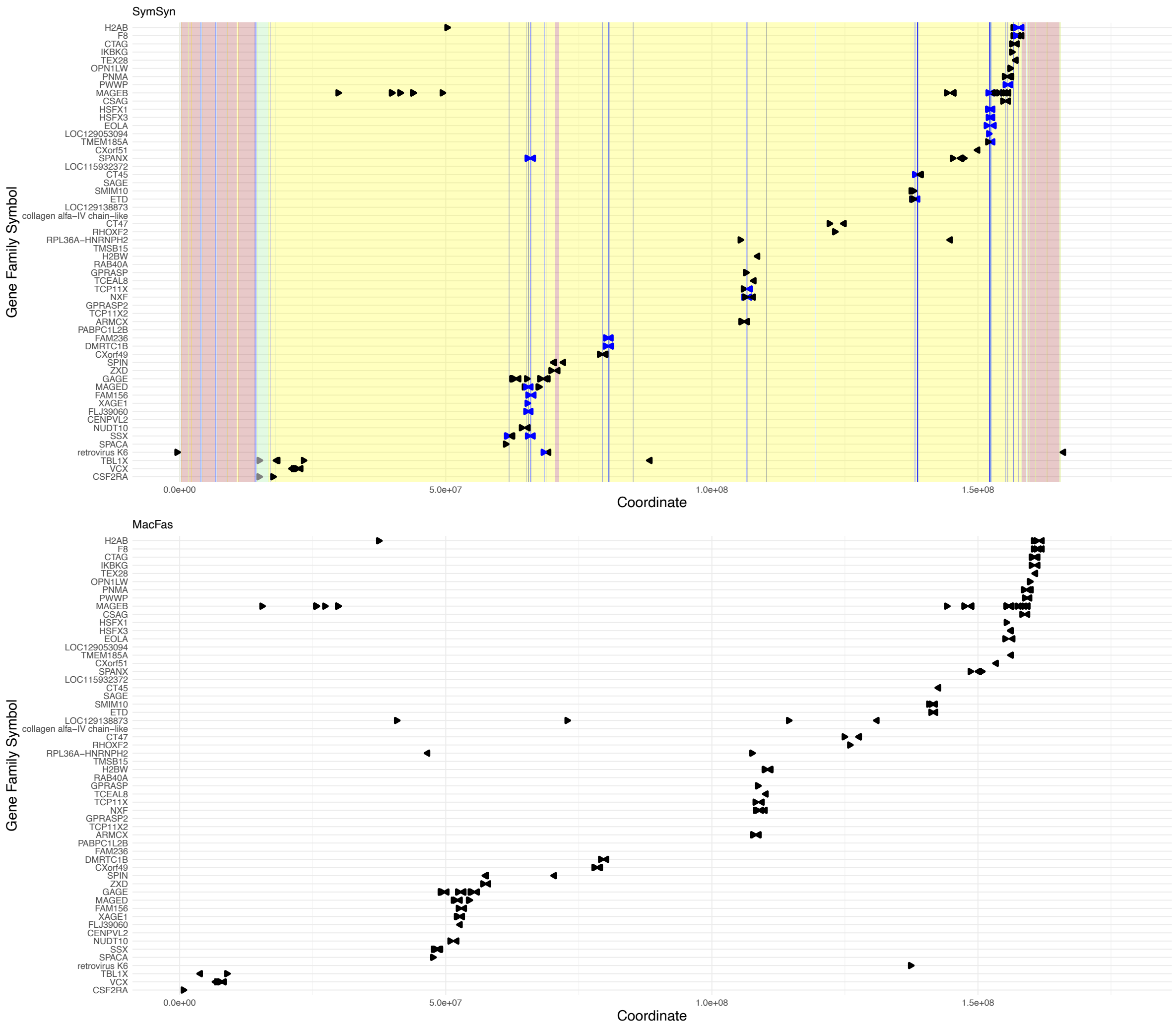


**Supplementary Fig 5. Gene family location and orientation on the X.** Each gene copy is represented as an arrow indicating strand orientation (right-pointing = forward strand, left-pointing = reverse strand). Gene families are ordered by the coordinates of their largest copy cluster. Copies within palindromes are coloured blue. Chromosome backgrounds indicate sequence classes as classified by Makova et al. 2024 (ampliconic, ancestral, palindrome, PAR, satellite; not available for macaque). Chromosome lengths are scaled to the longest X chromosome across species (gorilla). Species abbreviations: PanPan (bonobo), PanTro (chimpanzee), HomSap (human), GorGor (gorilla), PonAbe (Sumatran orangutan), PonPyg (Bornean orangutan), SymSyn (siamang), MacFas (macaque).

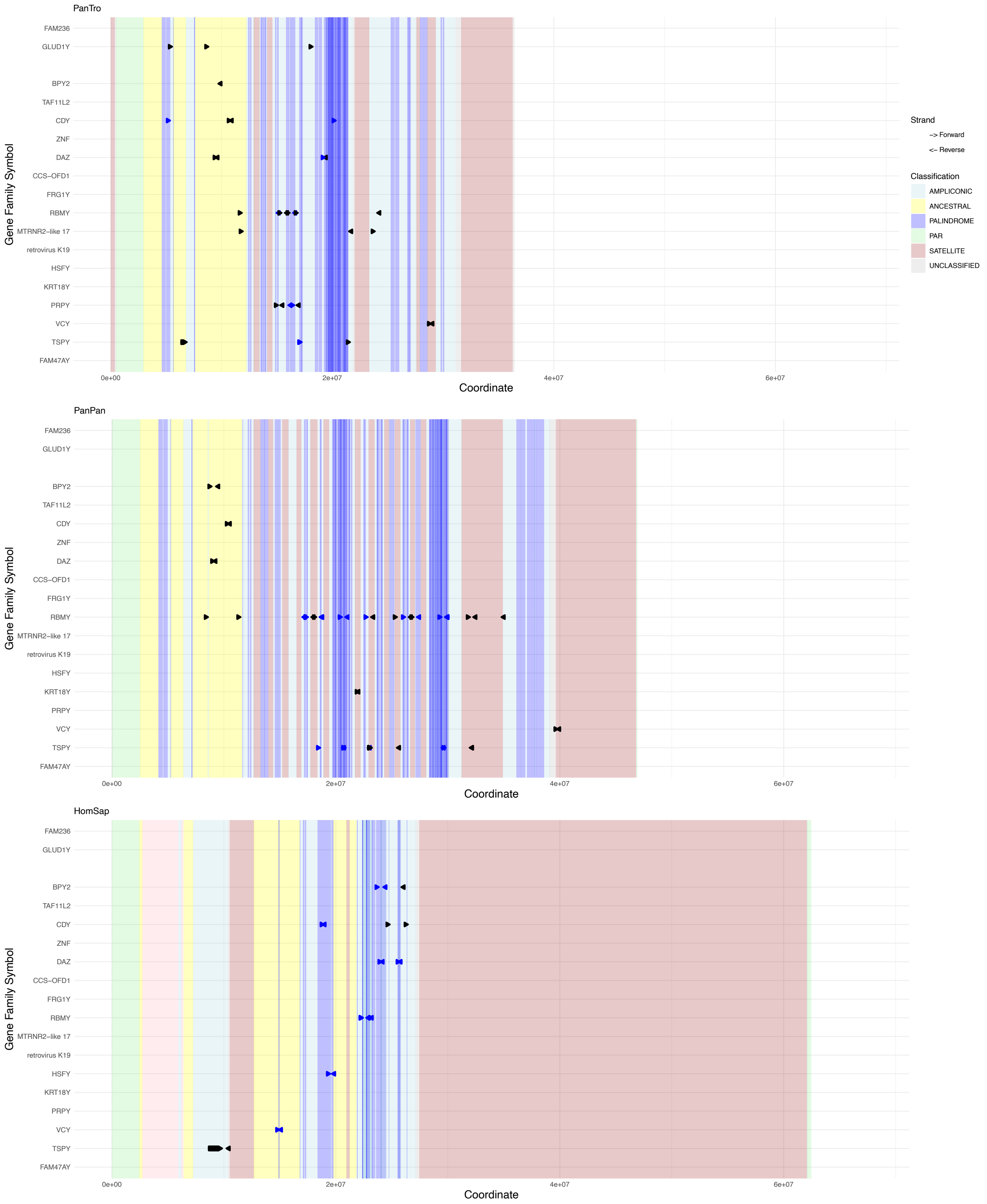

PanTro
PanPan
HomSap
FAM236
GLUD1Y
BPY2
TAF11L2
CDY
ZNF
DAZ
CCS−OFD1
FRG1Y
RBMY
MTRNR2−like 17
retrovirus K19
HSFY
KRT18Y
PRPY
VCY
TSPY
FAM47AY
Gene Family Symbol
0e+00
2e+07
4e+07
6e+07
Coordinate
Strand
−> Forward
<− Reverse
Classification
AMPLICONIC
ANCESTRAL
PALINDROME
PAR
SATELLITE
UNCLASSIFIED

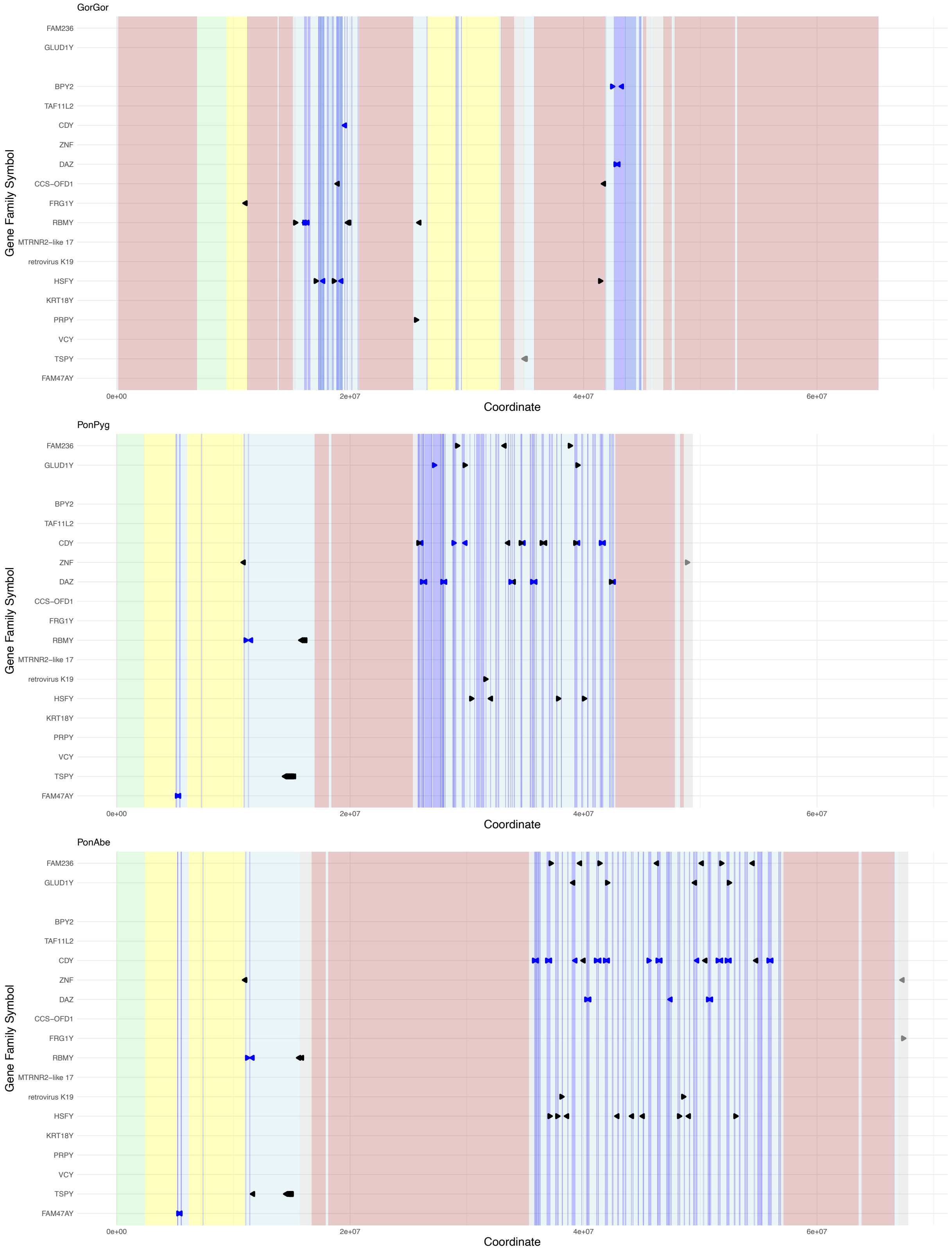
GorGor
PonPyg
PonAbe
FAM236
GLUD1Y
BPY2
TAF11L2
CDY
ZNF
DAZ
CCS−OFD1
FRG1Y
RBMY
MTRNR2−like 17
retrovirus K19
HSFY
KRT18Y
PRPY
VCY
TSPY
FAM47AY
Gene Family Symbol
0e+00
2e+07
4e+07
6e+07
Coordinate

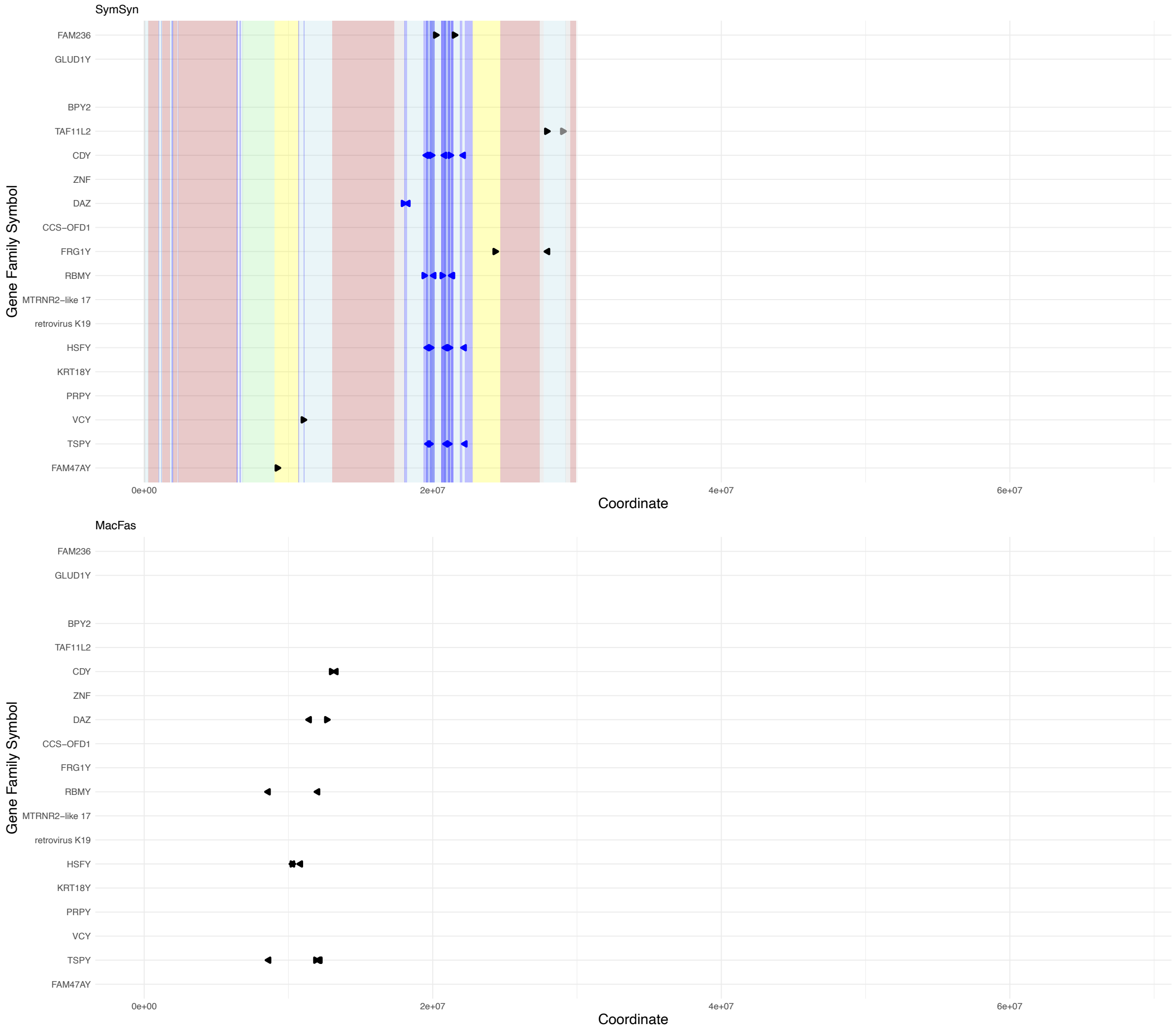


**Supplementary Fig 6. Gene family location and orientation on the Y.** Each gene copy is represented as an arrow indicating strand orientation (right-pointing = forward strand, left-pointing = reverse strand). Gene families are ordered by the coordinates of their largest copy cluster. Copies within palindromes are coloured blue. Chromosome backgrounds indicate sequence classes as classified by Makova et al. 2024 (ampliconic, ancestral, palindrome, PAR, satellite; not available for macaque). Chromosome lengths are scaled to the longest X chromosome across species (S. orangutan). Species abbreviations: PanPan (bonobo), PanTro (chimpanzee), HomSap (human), GorGor (gorilla), PonAbe (Sumatran orangutan), PonPyg (Bornean orangutan), SymSyn (siamang), MacFas (macaque).

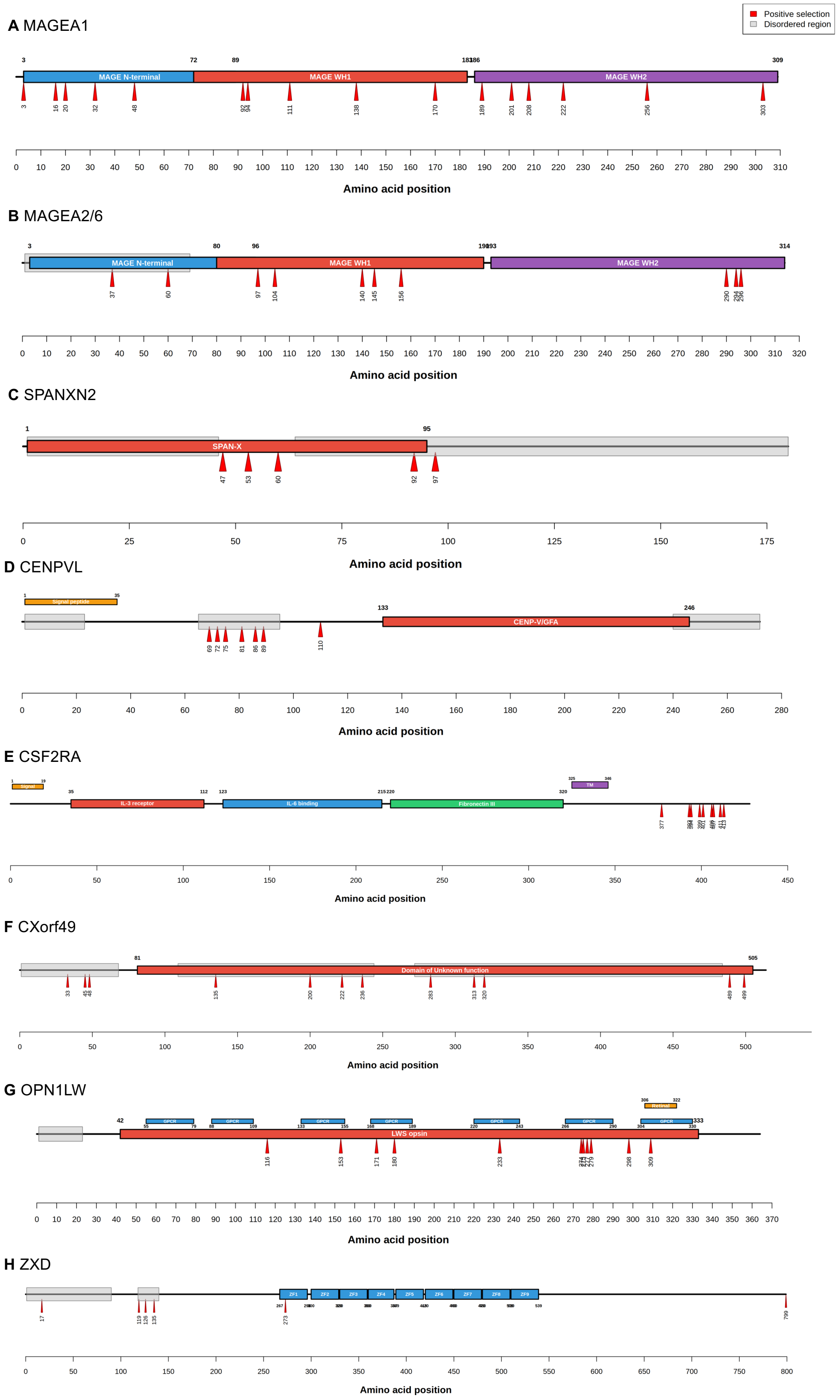


**Supplementary Figure 7. Distribution of positively selected sites across gene families**. Protein domains are based on human amino acid sequences as predicted by InterPro (Blum et al., 2025). Positively selected sites were identified by PAML site tests. Gene families are grouped by overall selection pattern **a-c**: Neutral (MAGEA1, MAGEA2/6, SPANXN2); **d-h:** Purifying with positively selected sites (CENPVL, CSF2RA, CXorf49, OPN1LW, ZXD).

### Supplementary Notes

**Supplementary Note 1: Positive selection in gene families under neutral or purifying selection.**

Interestingly, several gene families showed positively selected sites despite exhibiting overall neutral or purifying selection across the species tree. These families may harbour positively selected sites amid broader neutral or purifying selection.

Gene families showing overall neutral selection with positively selected sites (MAGEA1, MAGEA2/6, SPANXN2/3) displayed dispersed patterns similar to rapidly evolving families. In MAGEA2/6, positively selected sites were distributed throughout the protein sequence without clear hotspots, occurring in both the N-terminal region and both MAGE helices. MAGEA1 showed a comparable pattern (Supplementary Figure 7). In SPANXN2, four of five selected sites were in the SPAN-X domain shared by all SPANX family members.

Gene families evolving under purifying selection also showed positively selected sites. Some displayed dispersed patterns: OPN1LW showed positively selected sites distributed throughout the long-wave-sensitive opsin region, with four sites clustered in one G-protein-coupled receptor domain. CXorf49 showed positively selected sites distributed throughout the protein, which is almost entirely intrinsically disordered and lacks known domains.

Other purifying selection families showed clear hotspots of selected sites. In CENPVL, 6 of 7 positively selected sites clustered in a disordered region outside known domains. Similarly, all positively selected sites in CSF2RA clustered at the C-terminus outside known domains (Supplementary Figure 7). Most positively selected sites in ZXD occurred in disordered regions, with one site in the first zinc finger repeat.
